# Darwin’s finches - an adaptive radiation constructed from ancestral genetic modules

**DOI:** 10.1101/2021.09.17.460815

**Authors:** Carl-Johan Rubin, Erik D. Enbody, Mariya P. Dobreva, Arhat Abzhanov, Brian W. Davis, Sangeet Lamichhaney, Mats Pettersson, C. Grace Sprehn, Carlos A. Valle, Karla Vasco, Ola Wallerman, B. Rosemary Grant, Peter R. Grant, Leif Andersson

## Abstract

Recent adaptive radiations are models for investigating mechanisms contributing to the evolution of biodiversity. An unresolved question is the relative importance of new mutations, ancestral variants, and introgressive hybridization for phenotypic evolution and speciation. Here we address this issue using Darwin’s finches, which vary in size from an 8g warbler finch with a pointed beak to a 40g large ground finch with a massive blunt beak. We present a highly contiguous genome assembly for one of the species and investigate the genomic architecture underlying phenotypic diversity in the entire radiation. Admixture mapping for beak and body size in the small, medium and large ground finches revealed 28 loci showing strong genetic differentiation. These loci represent ancestral haplotype blocks with origins as old as the Darwin’s finch phylogeny (1-2 million years). Genes expressed in the developing beak are overrepresented in these genomic regions. Frequencies of allelic variants at the 28 loci covary with phenotypic similarities in body and beak size across the Darwin’s finch phylogeny. These ancestral haplotypes constitute genetic modules for selection, and act as key determinants of the exceptional phenotypic diversity of Darwin’s finches. Such ancestral haplotype blocks can be critical for how species adapt to environmental variability and change.

---

Identification of the factors that promote or constrain the process of adaptive radiation, or the proliferation of forms from a single common ancestor, provide opportunities for understanding the origins of biodiversity. Species that radiate rapidly are thought to share some common features (*1*), promoting their ability to evolve into diverse forms (*2, 3*), whereas depauperate clades may lack them (*4, 5*). One of these features, evolvability, may be determined in part by the modularity of phenotypic traits (*6*), allowing some species to exploit ecological opportunity more readily (*3*). Two factors that influence why some species exhibit greater evolvability than others are phenotypic plasticity and the genetic potential for diversification (*3*). While the rapid speciation in adaptive radiations provides limited time to generate *de novo* genetic variation, ancestral polymorphisms can facilitate rapid accumulation of diverse combinations of alleles (*7–12*). Under this model, ancestral variation is sorted in unique combinations in descendent lineages (*13*) and/or is transmitted across lineages through introgression (*9, 11*). Hybridization may lead to loss of genetic differentiation (*14, 15*), but may also enhance the potential for selection by increasing phenotypic and genetic variation (*16, 17*). The identification of genetic variants underlying phenotypic variation is essential for understanding the role of ancestral genetic variation in evolutionary change. This remains an outstanding challenge when comparing species in the absence of genetic data because causal variants are greatly outnumbered by neutral variants. However, recent adaptive radiations, in particular those that still hybridize, are excellent groups for studying the origins of genetic variation and their effect on phenotype because gene flow has homogenized the genetic background, thus facilitating the identification of loci contributing to phenotypic differences among species (*18, 19*).

The Darwin’s finch radiation comprises 18 species, 17 present in Galápagos and one on Cocos Island. The group is highly unusual in that no species is known to have become extinct as a result of human activities, in contrast to some other avian radiations (*20*). The species have experienced current and historical gene flow (*21–24*) and diversification involved a key ecological trait, beak morphology, that mediates efficient use of different food sources (insects, seeds of various sizes, cactus fruits, and even blood from other birds) (*25*). Previous genetic studies have revealed a few loci where ancestral alleles explain variation in beak morphology: *ALX1* affecting beak shape (*21*), a genomic region controlling beak size including *HMGA2* and three other genes (*MSRB3, LEMD3, WIF1*) (*26, 27*), and in addition a number of suggestive loci under selection (*21, 26–28*). Whether other loci mediating phenotypic evolution in this group also represent ancestral variation and their role in phenotypic evolution in the radiation is unknown. Here, we present a high-quality chromosome-scale reference genome and leverage a natural scaling transformation in beak size (*29*) across three species of ground finches (*Geospiza*) to identify 28 loci under selection. We show that these haplotype blocks linked to phenotypic divergence are as old as the Darwin’s finch phylogeny. These genetic modules have been reused over the last million years, were exchanged by gene flow, and contributed to the rapid phenotypic evolution and speciation among Darwin’s finches.

## High quality assembly of the *Camarhynchus parvulus* genome

The previously reported genome assembly based on Illumina short reads of a medium ground finch (*G. fortis*) is highly fragmented (*30*). We therefore decided to develop a high quality, highly contiguous assembly for Darwin’s finches by combining long-read data with chromatin contact (HiC) data (**Fig. S1**). Because of the uncertainty to export tissue samples with intact long DNA molecules from the Galápagos National Park we carried out Oxford Nanopore Technologies (ONT) sequencing at the Galápagos Science Center. Genomic DNA prepared from a male small tree finch (*C. parvulus)* was of high molecular weight and this individual was selected for Oxford Nanopore Technologies (ONT) sequencing. The close evolutionary relationship among all species of Darwin’s finches (pairwise interspecies *dXY* in the range 0.2-0.3% (*31*)) implies that this reference assembly can be used across the phylogeny. We generated 35X ONT long read sequence coverage from the reference individual **(Fig. S1**). We corrected erroneous base calls using linked-read data and generated chromosome-size scaffolds using HiC data. The resulting assembly is of similar quality to current state-of-the art genome assemblies in contiguity and accuracy (96% of the sequence assigned to chromosomes, N50 = 71.1 Mb, BUSCO = 96.1% complete, gaps = 0.01%) and shows a high degree of conserved synteny to the zebra finch genome assembly (**Fig. 1a**). Field-collected tissue samples were used to generate RNA-seq data for annotation (**Table S1**; **Fig. S1**). Genome annotation of these data using the Ensembl annotation pipeline (*32*) generated 17,167 gene modules that include non-coding RNA and microRNA. We further used 25 *C. parvulus* individuals to generate a linkage-disequilibrium based recombination map (*33*) (**Fig. S2**). Consistent with other avian species (*34, 35*), recombination rates are generally elevated at the ends of chromosomes and correlated with nucleotide diversity (*R*^*2*^ = 0.19, *P* < 0.001), particularly on chromosome Z (*R*^*2*^ = 0.27, *P* < 0.001). This relationship is consistent with the widespread effects of background selection known in birds (*35, 36*).

**Fig. 1.**
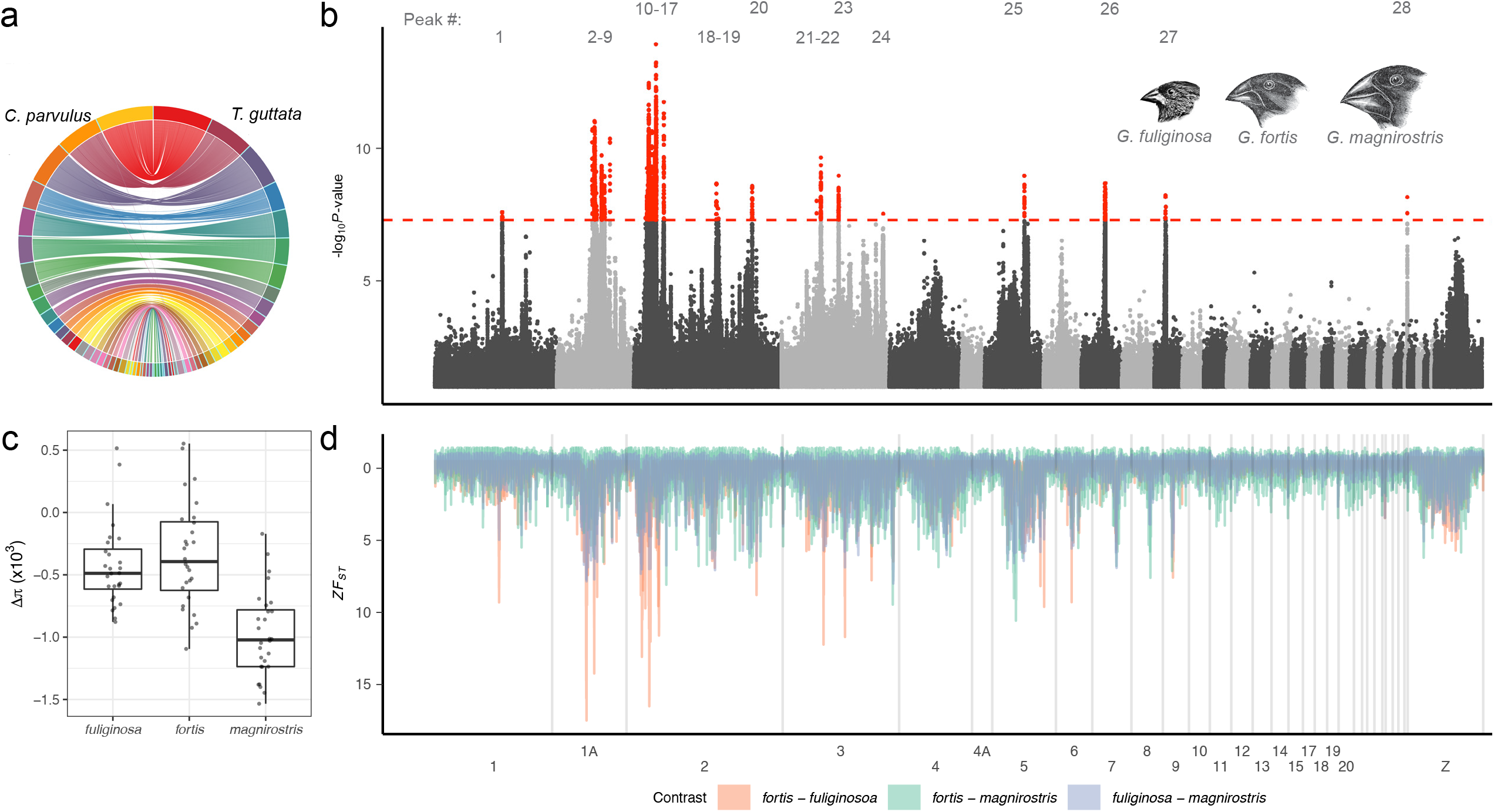
Genome assembly and genetic differentiation among three species of Darwin’s finches. (**a**) Illustration of conserved synteny to zebra finch (*T. guttata*). (**b**) Genome-wide admixture mapping using three species sorted ascendingly by beak and body size: 0 = *fuliginosa*, 1 = *fortis* and 2 = *magnirostris*. The dotted red line indicates the significance threshold set by permutation. Illustrations of the three species are adapted from P.R Grant and Darwin(*56*). (**c**) Boxplot showing the difference in nucleotide diversity between the regions of association marked in (b) and regions outside the main area of association. Centreline indicates the median, bounded by the 25^th^ and 75^th^ percentile, with whiskers extending to 1.5x the interquartile range. (**d**) Genome-wide *F*_*ST*_ for all three possible pairwise combinations of *Geospiza* considered here. Lines are coloured by the comparison of interest. Many highly divergent regions are shared between contrasts and overlap with regions of association in (b) (**Data S2** and S**3**).

## Twenty-eight trait loci explaining phenotypic differentiation

We used admixture mapping (*37*) to search for loci contributing to genetic differentiation between three closely related species that differ primarily in a scaling factor for beak and body size from small to large (*29*): the small, medium and large ground finches (*G. fuliginosa*, *G. fortis* and *G. magnirostris*, respectively) (**Fig. S3**). This trio was selected based on their striking phenotypic differentiation in beak and body traits alone and low genome-wide genetic differentiation (pairwise *F*_*ST*_ = 0.02 – 0.10), because this reduces the background noise due to genetic drift. In this study, we generated whole genome, short read sequence data from 28 individuals from these three species and combined these with previously published samples for a total of 75 birds on 9 islands (mean coverage = 17 ± 9, **Table S2; Data S1**) and applied phenotypic scores of 0,1,2 to reflect the increasing beak and body size of *G. fuliginosa < G. fortis < G. magnirostris,* because phenotypic data were not available for each individual. The experimental setup is similar to an earlier study using these three species, but this studied used only one island and reduced representation sequencing (*27*). Admixture mapping (*37*) revealed 28 loci that exceeded the significance threshold set by permutation (**Fig. 1b**) and represent independent loci (**Data S2**). The size of these regions ranges from thousands of kb to 2.7 Mb (**Fig. 2**) and contained between 0 and 35 genes (**Data S2)**. Outlier loci were clustered on macrochromosomes and included the previously described *ALX1* and *HMGA2* loci affecting beak morphology, both located on chromosome 1A and only ~7 Mb apart (locus 4 and 9 in **Fig. 1b**). These loci are separated by a recombination hot-spot (**Fig. S2**), consistent with previous results that these loci do not show strong linkage disequilibrium (*21, 26*).

**Fig. 2.**
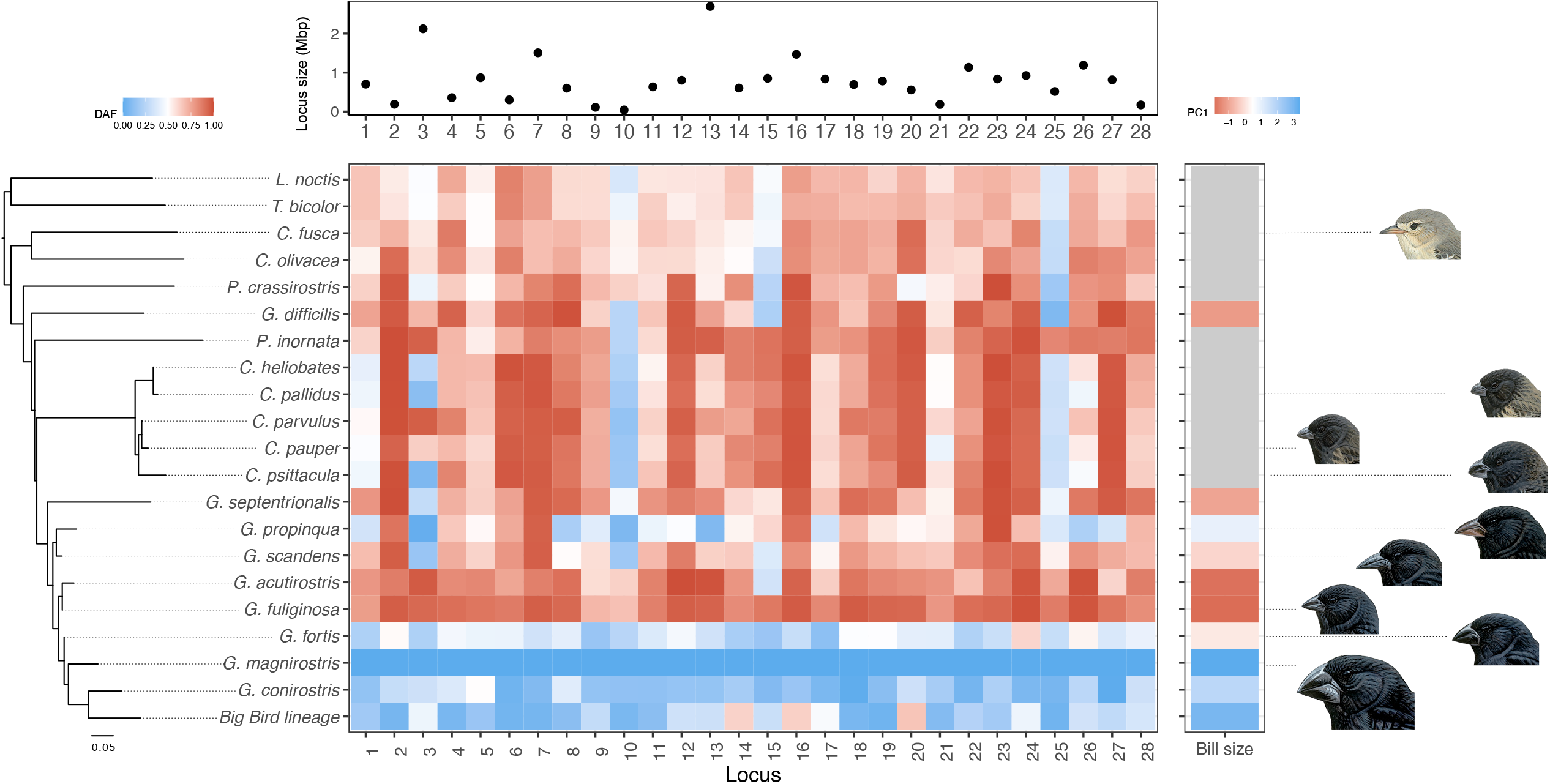
Haplotype variation across the Darwin’s finch phylogeny. The heatmap displays average delta allele frequency (DAF) based on 34–328 SNPs/locus for each species compared to *G. magnirostris*, the species with the largest beak. On the right, bill size is presented according to a principal component analysis of three beak dimensions averaged across all island populations for each species. Only *Geospiza* species are shown. Above, the size of each genomic region (in Mb) is marked in a dot plot. Right, finch illustrations reproduced by permission of Lynx Edicions.

Regions of association identified with admixture mapping largely mirrored the results of *F*_*ST*_-based contrasts (**Fig. 1d**) and strongly correlated with per-window estimates in the two contrasts involving *G. fuliginosa (R*^*2*^_*fortis-fuliginosa*_ = 0.84, *R*^*2*^_*magnirostris-fuliginosa*_ = 0.69, **Data S2**). We do not expect a perfect match between the results of admixture mapping and *F*_*ST*_, analysis because the former is based on a linear comparison of the trio while the latter is derived from pairwise comparisons between species. In the 28 regions of association, *G. fuliginosa* and *G. magnirostris* were often homozygous for different haplotypes while *G. fortis* exhibited intermediate allele frequencies (**Fig. S4**; **Data S3**). This is highlighted at the *HMGA2* locus on chromosome 1A, where measures of Tajima’s D are strongly negative for *G. fuliginosa* and *G. magnirostris,* but strongly positive for *G. fortis* (**Fig. S4**), consistent with balancing selection maintaining haplotype diversity in the phenotypically variable *G. fortis* population(*25*). Nucleotide diversity was reduced in *G. magnirostris* relative to the genomic background in 23 of the 28 regions (**Fig. 1c**), consistent with selective sweeps in *G. magnirostris* or an ancestor. These regions also fell in genome-wide low-recombination regions (mean ρ in peaks = 1.4 compared with mean ρ outside = 2.0), with the exception of one peak on chromosome 25 (ρ = 4.7), consistent with previous studies in Darwin’s finches that regions of elevated differentiation often lie in recombination cold spots (*31*). Low recombination in these regions has likely facilitated the persistence of large haplotype blocks despite high gene flow amongst Darwin’s finch species, as predicted from theoretical studies (*36*).

To explore the extent to which the loci detected in the ground finch contrast also shape phenotypic diversity among tree finches (*Camarhynchus*), we next performed a similar admixture analysis comparing small, medium and large tree finches (*C. parvulus*, *C. pauper* and *C. psittacula*, respectively), also classified as 0,1,2, respectively, based on beak and body size. These samples were previously sequenced and include 46 individuals (n = 10-25 each) from 8 islands. This replicated a signal for the *HMGA2* locus affecting beak size (*P*=4×10^−16^) (**Fig. S5**), as expected from previous work (*26*). No other locus showed such a striking signal of genetic differentiation, but we noted an overlap of higher genetic differentiation, approaching genome-wide significance, for several of the loci detected in the ground finch contrast (**Fig. S5**). Nevertheless, the identification of regions associated with phenotypic variation in size among *Camarhynchus* may be hampered by our comparatively smaller sample size than in *Geospiza* (n = 75 vs. n = 46).

In order to determine the evolutionary origin of haplotypes at the 28 loci detected in the ground finch contrast, we compared allele frequencies at the most differentiated SNPs across all 18 species of Darwin’s finches, two outgroups *L. noctis* and *T. bicolor.* We also included the Big Bird hybrid lineage, which was formed by the mating of a *G. conirostris* male and two *G. fortis* females(*24*). These included genome sequences from previously published data and 62 new individuals (*n* = 321 in total, **Data S1**). If genetic differentiation among *Geospiza* is caused by *de novo* mutations that occurred after the split from *Camarhynchus*, we would expect to find little shared haplotype structure in non-*Geospiza* species as a consequence of random accumulation of variants. In sharp contrast, the allele frequency comparison revealed a non-random pattern for the most differentiated SNPs at the 28 loci (**Fig. 2**). Notably, a few *G. magnirostris* haplotypes are present at a relatively high frequency across the radiation (e.g., 10, 25), while most are consistently highly differentiated from haplotypes in other species except those with relatively large beaks. Furthermore, a large portion of *G. magnirostris* major alleles, 67% (1,273/1,914), were derived relative to outgroups *L. noctis* and *T. bicolor* consistent with selective sweeps in the *G. magnirostris* lineage (**Data S4**). Together, these results imply that the “large” and “small” haplotype blocks identified by admixture mapping in *Geospiza* predates the separation of *Geospiza* and *Camarhynchus*. Heterogenous combinations of these haplotypes in two species with large beaks, *G. propinqua* and the Big Bird lineage (**Fig. 2**), raise the intriguing possibility that unique combinations of alleles are formed during the speciation process by incomplete lineage sorting (ILS) and/or introgression, and retained by natural selection.

We evaluated the hypothesis that these haplotype blocks are old by first generating a neighbor-joining genetic distance tree for the entire radiation by concatenating the most differentiated SNPs from each genomic region. The topology of this tree matches previous species-tree reconstructions using other phylogenetic models (*21, 24*). We predicted that, if these haplotypes carry causative mutations for species phenotypes, then phenotypically similar species will cluster together in this “haplotype tree”. We compared the topology of this tree with a species tree based on 4.9 million high-quality SNPs from the rest of the genome and found them to be discordant (**Fig. 3a**); individual trees for each of the 28 loci are given in **Fig. S6**. We further analysed whether allele frequencies of differentiated SNPs at each of the 28 loci were more concordant with the haplotype tree or with the species tree and concluded that, for all loci except two, allele frequencies were more concordant with the haplotype tree (**Fig. S7**). In the species tree, the earliest divergence involves the warbler finches and all other finches in the radiation (**Fig. 3a**). In contrast, in the haplotype tree, the most divergent grouping separates the largest ground finches (*G. magnirostris, G. fortis* and *G. conirostris*) from all other species. Further, *G. fuliginosa* and *G. acutirostris* cluster with *Pinaroloxias inornata* (the Cocos Island finch), which all share a common small phenotype (small bodies and small pointed beaks), but are not monophyletic on the species tree. At all 28 loci, except one (#10), the haplotypes present in *P. inornata* are more similar to those in *G. fuliginosa* than in *G. magnirostris* (**Fig. 2**). This implies that the finch that colonized Cocos Island carried the small variant alleles at most of these 28 loci and subsequently diverged through local adaptation and genetic drift in geographical isolation from other ancestral finch populations. In contrast, the small and medium ground finches, two sympatric species that differ in feeding ecologies and exchange genes (*25*), are essentially indistinguishable genetically, except at the 28 loci under selection (**Fig. S3b**). Finally, we confirmed deep divergence (mean age 1.3 ± 0.5 MYA) of the major haplotypes in *G. fuliginosa* and *G. magnirostris* by estimating the time to the most recent common ancestor at each locus using the net frequency of nucleotide substitutions (*d*_*a*_) and the estimated substitution rates in Darwin’s finches (*21*) (**Fig. 3b**).

**Fig. 3:**
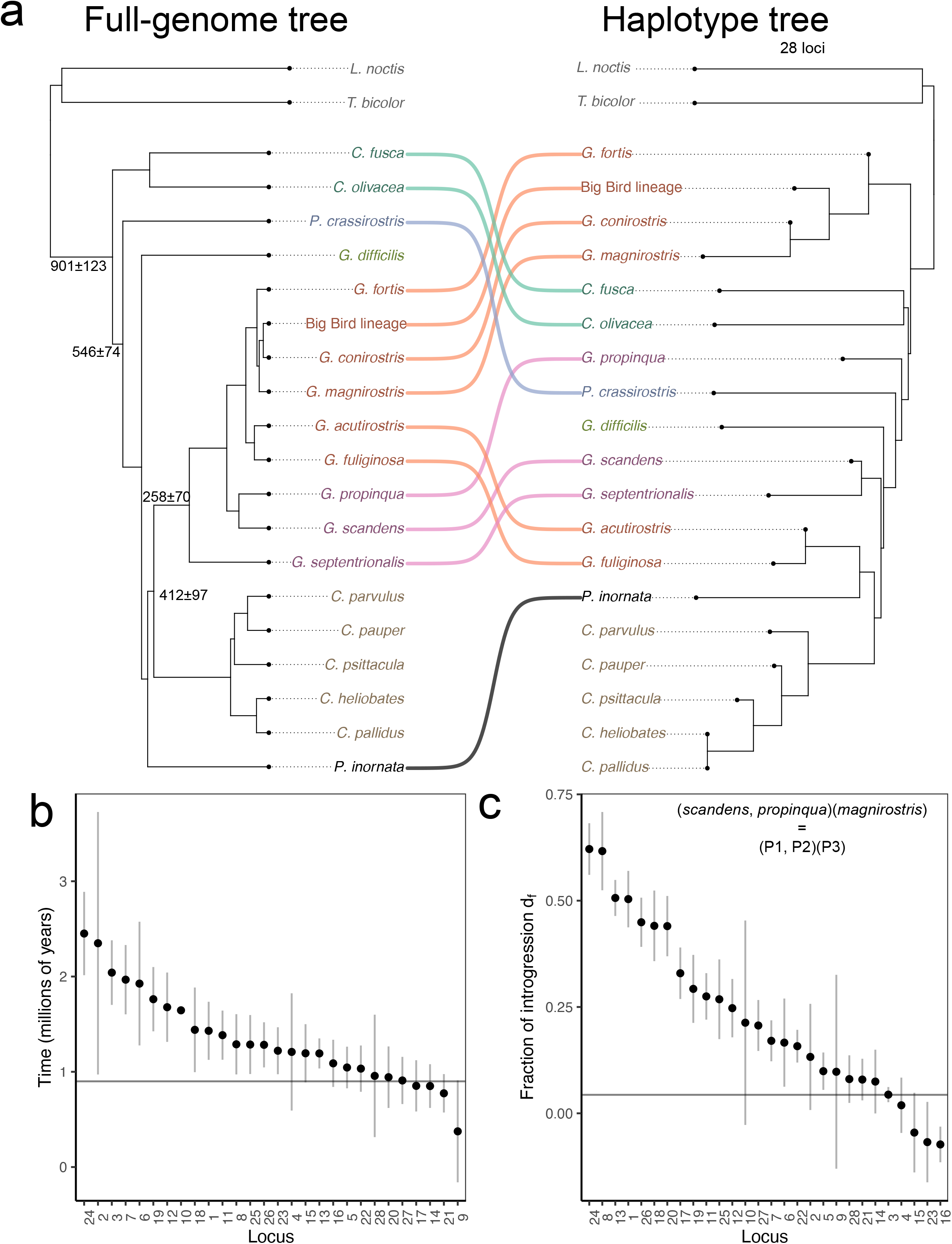
Characterization of 28 adaptive loci. **(a)** Left, a neighbor-joining tree for all species of Darwin’s finches based on 4.9 million SNPs. The tree was converted to a chronogram using ape(*57*) and branching times are reproduced from Lamichhaney *et al*.(*21*). Right, a neighbor-joining tree for the 28 concatenated loci identified in the association analysis. Species names are coloured by *a-priori* clade assignments and a co-phylo diagram(*58*) highlights changes in topology between the trees. **(b)** Time to most recent common ancestor between *G. fuliginosa* and *G. magnirostris* haplotypes for all 28 loci. Time was estimated by the conversion of genetic divergence (*d*_a_) to time using *T* = *d*_a_/(2μ) and a mutation rate of 2.04 × 10^−9^ (Ref. (*21*)). The horizontal line marks 900,000 years, the approximate time of divergence between warbler finches and all other finches in the radiation(*21*). **(c)** Fraction of introgression from *G. magnirostris* to *G. propinqua* on Genovesa island for each of the 28 loci, as measured by *d*_*f*_. Loci are arranged by the strength of the introgression and a line is drawn at the median of genome-wide *d*_*f*_ = 0.04. The trio arrangement is written on the graph and ABBA/BABA statistics listed under methods. In (b) and (c) error bars represent 95% confidence intervals.

## Introgression of ancestral haplotypes

The presence of distinct combinations of haplotypes across the phylogeny indicates ILS or introgression. The Genovesa cactus finch *G. propinqua* has the fourth largest beak of all *Geospiza*, but the pointed beak characteristic of other cactus finches, and carries a mix of large and small haplotypes (**Fig. 2**). Gene flow from *G. magnirostris* to *G. propinqua* has been implicated from field observations (*16*). We estimated the fraction of introgression from *G. magnirostris* to *G. propinqua* on Genovesa using *df*, which incorporates *dXY* into an extension of ABBA/BABA D statistics (*38*), and found that regions of high *G. magnirostris* similarity share an excess of derived alleles (**Fig. 3c),**and often reduced genetic divergence *dXY* (**Data S5)**, consistent with introgression. The role of gene flow in generating distinct combinations of haplotypes is also evident in the Big Bird lineage (**Fig. 2**). The Big Bird lineage is characterized by an unusually large beak on a relatively small body (*24*). The combination of *G. conirostris*, a sister species to *G. magnirostris*, and *G. fortis* alleles resulted in a unique phenotype and genotype, with most loci sharing greater affinities with *G. conirostris* than with *G. fortis* (**Fig. 2**). It remains an outstanding challenge to distinguish between ILS and introgression using D statistics (*39*), but our results and field observations strongly imply that gene flow can transfer these haplotypes among species, as previously demonstrated for *ALX1 (22).*

## Trait loci are enriched for developmental genes

Ancient haplotypes may be retained in descendent species even if they are neutral. However, the loci identified here segregate with phenotypic variation among the species and are expected to contain alleles important for beak and body size variation. We conducted an enrichment analysis using mouse orthologs for the genes in the vicinity of the 28 loci using the software GREAT (*40*) and found that deleterious mutations at these loci were significantly associated with abnormal development of cartilage and bones (**Fig. 4a; Fig. S8**). Furthermore, these mouse orthologs were significantly enriched for genes expressed during craniofacial and limb development, which is consistent with our expectation that genetic changes at many of these 28 loci affect beak development. The enrichment includes genes spread across the 28 loci (**Data S2**).

**Fig. 4.**
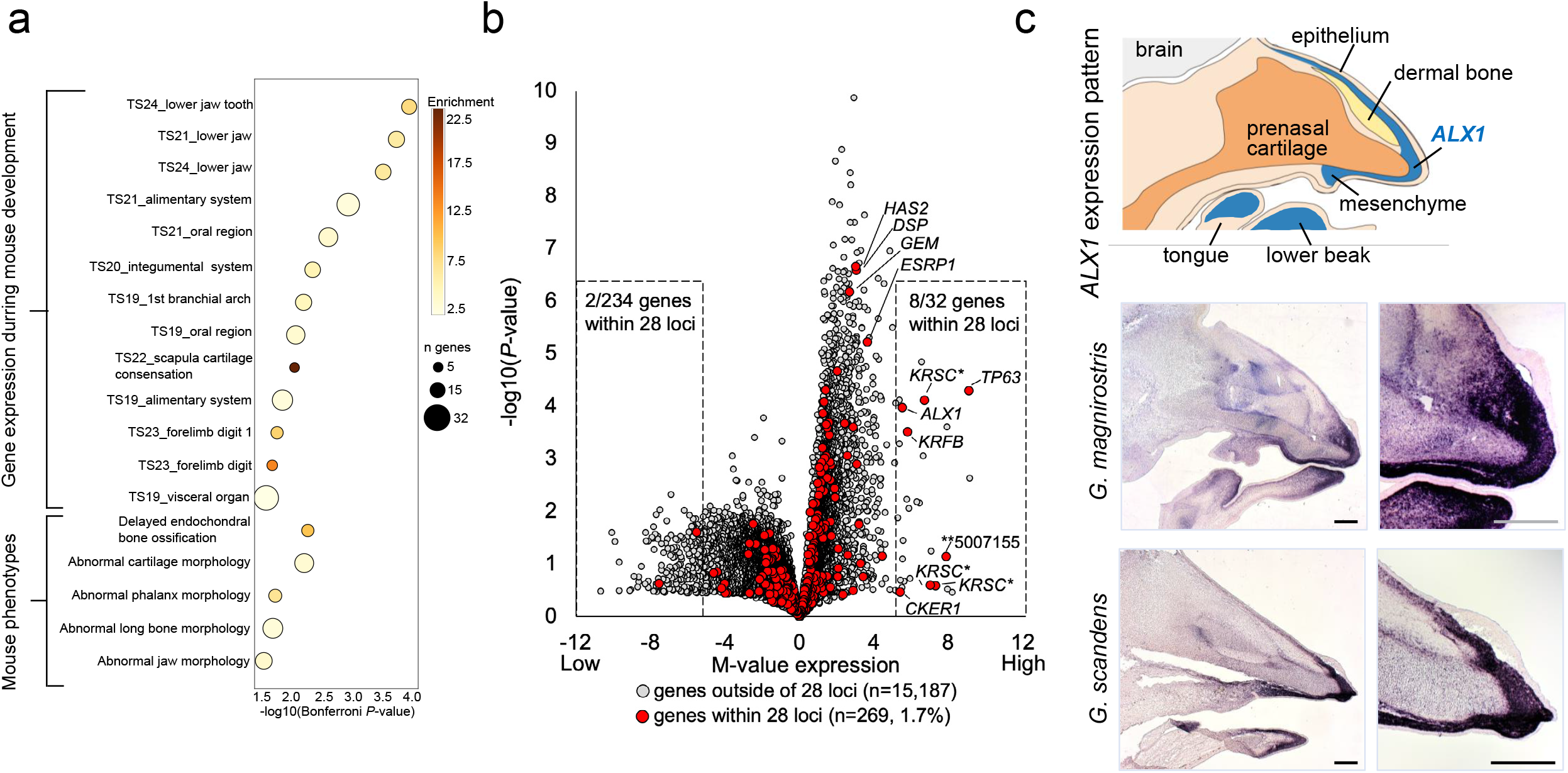
Enrichment analysis and gene expression. **(a)** Annotation term enrichment analysis. GREAT(*40*) was used to screen for enrichment of gene annotation terms associated with the 28 differentiated regions. −log10 Bonferroni corrected *P*-values of significantly enriched terms are shown on the x-axis. Fold enrichment is indicated using dot colours. Dot sizes indicate numbers of genes belonging to each annotation term. **(b)** Gene expression in Darwin’s finch upper beak vs. other tissues. RNAseq gene expression levels in upper beak (n samples=9) were compared with expression levels in non-craniofacial tissues (n samples=7, n tissues=4). −log10 *P*-values for differential expression are shown on the y-axis and M-values (log2(fold change beak samples vs. other tissues)) are shown on the x-axis. Dotted boxes show genes with M-values <−5 and >+5, representing genes with lower (left) and higher (right) expression in beak samples (right) compared with other tissues. The asymmetry in −log10(*P*-values) is a result of comparing one tissue (beak) to several other tissues. Gene names are shown for selected genes. * Separate genes belonging to a gene cluster of scale keratins. ** is a gene of uncertain function (see **Data S6). (c)** *In situ* hybridization. Top: schematic representation of the expression pattern of *ALX1* (blue) in E7 embryos of zebra finch (n=5) and Darwin’s finches (9 species, n=21). Bottom: mRNA expression of *ALX1* (dark purple) in mid-face longitudinal sections through the heads of Darwin’s finch embryos. The developing beak region is shown. Magnified area of the left images is shown on the right. Scale bar: 250 μm.

Because of the difficulties in interpreting gene-by-term enrichment data (*41*), we performed two types of analyses to validate gene expression. RNA-seq of upper beaks from 9 embryonic day 7 (E7) Darwin’s finch embryos representing 6 species confirmed that a number of the genes within the 28 loci are expressed in the developing beak and these loci were 14-fold enriched for genes with higher expression levels (M≥5 = fold change ≥32) in the developing beak compared with other tissues (*χ*^2^-test, *P*=7.4×10^−24^, d.f.=1; **Fig. 4b**). We carried out *in situ* hybridization (ISH) of two candidates for craniofacial development (*ALX1-* locus 7 and *RUNX2*-locus 24). ISH data on a total of 7 zebra finch (*Taeniopygia gutatta*) and 27 Darwin’s finch embryos (of 9 species) revealed that *ALX1* expression was strongly biased to the beak region over the developmental period when beak size and shape are established (E6-E7)(*42*), **Fig. 4c; Figs S9, S10a**). Additionally, we confirmed a similar expression pattern in zebra finch embryos for *RUNX2* (alias *OSF2*) *–* an essential gene for ossification of the mesenchyme (*43*) (**Fig. S10b**) located within a strong signal of differentiation among *Geospiza* (**Fig. S4b**).

## Discussion

We have identified ancestral haplotypes at 28 loci that have evolved by natural selection, shaping phenotypic diversity among Darwin’s finches throughout their evolution. Our functional characterization of these loci contributes to a growing body of literature suggesting that genetic differences between species of Darwin’s finches are enriched for genes involved in the key pathways for growth and beak development (*21, 26–28, 42*). Together these results extend the key role for the beak of the finch in ecological adaptation in this group. In this study, we show that the distribution of these genetic modules across the phylogeny reflects natural selection and most likely both incomplete lineage sorting and introgression (*21, 22*). Importantly, it is not only the presence/absence of these haplotype blocks that affects the phenotype but their frequency within species, illustrated by *G. fortis* that have intermediate haplotype frequencies at many of these loci (**Fig. 2; Data S4**). Intermediate frequencies, indicative of balanced polymorphism, provide the underlying variation for selection to sort adaptive haplotypes during speciation. Our findings support previous suggestions that ancestral variants contribute to phenotypic diversity, as indicated for pigmentation phenotypes among other songbird species (*8, 13*), colour morphs in the common wall lizard (*44*), colour patterns in *Heliconius* butterflies (*11*), various phenotypic traits in cichlids (*7, 9*), craniofacial morphology in pupfish (*45*), winter coat in snow-shoe hares (*46*), and adaptation to high altitude in humans (*47*). That ancestral variation can be retained in large populations preceding speciation is illustrated in Atlantic herring and stickleback where ecotypes show differences in the frequency of haplotype blocks at hundreds of loci, all underlying ecological adaption (*48, 49*).

Characteristic features of these 28 loci are the large size of the haplotype blocks, often spanning hundreds of kilobases, and their ancient origins, 1-2 million years ago (**Fig. 3b**). This is in contrast to another ancestral polymorphism in Darwin’s finches, at the *BCO2* locus controlling nestling beak colour, where a single base change constitutes the likely causal mutation in the absence of haplotype structure (*50*). The identification of causal variants within the haplotype blocks described here is challenging because of strong linkage disequilibrium among many sequence variants within each region. Such large haplotypes could include structural variants, which have been proposed as a key determinant of adaptive evolution and speciation (*51, 52*), but these 28 loci do not represent large inversions (>5 Mb) and tended to be relatively small (0.1-2.7 Mb) compared to e.g., supergenes (*51, 53*). Furthermore, none of the loci exhibit the sharp borders in our association analysis that is characteristic of inversions maintained as balanced polymorphisms (*48, 53*).

The block structures at the 28 loci most likely reflect large-effect haplotypes composed of clusters of multiple causal variants that have accumulated during the evolution of Darwin’s finches (*54*), similar to the evolution of alleles in domestic animals by the sequential accumulation of causal mutations during the last 10,000 years (*55*). The occurrence of these haplotype blocks in low recombination regions likely facilitated their evolution (**Data S2).** The reuse of ancestral genetic modules is a much faster route to adaptive change than the slow accumulation of adaptive *de novo* mutations (*7*). Our study is comprehensive in surveying genomic variation across all 18 extant species of a single adaptive radiation, and yet the principle finding, of repeated reassembly of ancient haplotype blocks in the formation of species, is likely to be a general feature of rapid radiations (*8, 45*) and of general importance for how species adapt to environmental variability and change (*25*).

## Author contributions

P.R.G. and B.R.G. collected the blood samples. L.A., C-J.R., P.R.G., B.R.G. and E.D.E. conceived the study. C.J.R, S.L. and K.V. performed ONT sequencing on Galápagos. C.J.R. and E.D.E. were responsible for the bioinformatic analysis. A.A. and M.P.D. collected embryonic material, prepared RNA samples and performed ISH. B.W.D. and M.P. contributed to the bioinformatic analysis. C.G.S. and O.W. contributed to experimental work. C.A.V. contributed to sample collection. L.A., E.D.E., B.R.G., P.R.G, and C-J.R. wrote the paper with input from other authors. All authors approved the manuscript before submission.

## Data availability statement

The ONT reads and the Illumina reads have been submitted to the short reads archive (http://www.ncbi.nlm.nih.gov/sra) under BioProject PRJNA743742.

## Code availability statement

The analyses of data have been carried out with publicly available software and all are cited in the Methods section. Code associated with bioinformatic analyses are available at: https://github.com/erikenbody/Darwins_finch_comparative_genomics.

## Acknowledgements

We thank Ashley Sendell-Price for helpful discussion on the manuscript and Fan Han for initial bioinformatic analysis. The collection of blood samples, funded by National Science Foundation (NSF), was conducted with annual permits from the Galápagos National Parks Directorate, with approval of Princeton University’s Animal Care Committee and in accordance with its protocols, and supported logistically by the Charles Darwin Research Station in Galápagos. We thank Oxford Nanopore Technologies for lending us sequencing equipment and for technical assistance related to library preparation and sequencing. Denye Ogeh and Fergal Martin at EMBL-EBI performed gene annotation. Blood samples were collected on San Cristóbal Island under permit number MAE-DNB-CM-2016-0041 to C.A.V., for which we thank Ministerio del Ambiente de Ecuador. Logistical support for researchers to enter the Galápagos and perform laboratory work was provided by GSC-Universidad San Francisco Quito. Tissue samples for expression analysis were collected with permissions PC-08-13 and PC-34-14 from Galápagos National Park; and MAE-DNB-CM-2016-0043 from Ministerio del Ambiente de Ecuador. The project was financially supported by Vetenskapsrådet and Knut and Alice Wallenberg Foundation. The National Genomics Infrastructure (NGI)/Uppsala Genome Center provided service in massive parallel sequencing and the computational infrastructure was provided by the Swedish National Infrastructure for Computing (SNIC) at UPPMAX partially funded by the Swedish Research Council through grant agreement no. 2018-05973.

## Author Information

The authors declare no competing financial interests. Correspondence and requests for materials should be addressed to L.A.

