## supplemental material for "Darwin’s finches - an adaptive radiation constructed from ancestral genetic modules"

#### Supplementary Methods

**Fig. S1**. **Genome assembly**. (**a**) Workflow for construction of the Camarhynchus_parvulus_V1.1 assembly. (**b**) Summary table of the assembly. (**c**) Tissues used for preparing RNA-sequencing libraries for genome annotation. (**d**) Summary table of the genome annotation (**e**) Circular plot visualizing the repeat content of the genome assembly. Chromosomes, scaled by size, are shown along the outside of the circle. LINE=Long Interspersed Nuclear Element, SINE=Short Interspersed Nuclear Element, LTR=Long Terminal Repeat, LCR=Low Complexity Repeat, SR=Simple Repeat.

**Fig. S2. Recombination map**. Recombination fraction estimated from linkage disequilibrium data for 25 *C. parvulus* samples. Population scaled recombination rate (ρ) was estimated using LDhelmet and ρ converted to cM/Mb using the conversion factor $r = \rho/2cN_e$ (*1*) and Ne = 33,861 (*2*). The inset, marked by green, highlights a recombination hotspot between the loci containing *ALX1* and *HMGA2* and $ZF_{ST}$ scores for the comparison of *G. fuliginosa*, *G. fortis* and *G. magnirostris* (see **Data S2**).

**Fig. S3**. **Genome-wide variation in three species of Darwin's finches.** (**a**) Maximum likelihood tree of concatenated genome-wide SNPs for all *G. fuliginosa*, *G. fortis* and *G. magnirostris* samples used in this study. (**b**) The first principal component of genome-wide variation is shown on the x-axis and on the y-axis is the first principal component using only the regions of associations identified in **Fig. 1b**. Incomplete separation can be seen using genome-wide SNPs, while the three species show separation using the regions of association. Points are coloured by species and each point is an individual, which were collected from a variety of islands (**Data S1**) **(c)** Average beak length and average beak depth for all *Geospiza* species and island combination listed in **Data S1** and used in comparative genomics analysis. Each point is coloured by its species designation, and there are multiple points for species we have sequenced on multiple islands. Selected species discussed in the manuscript are labelled. Note the four species with the greatest beak depth, also highlighted in **Fig. 2**, and the extensive phenotypic variation found in *G. fortis*.

**Fig. S4**. **Population genomic summary statistics for selected regions of association identified using admixture mapping.** (**a**) Peak 3 on chr1A containing *HMGA2*. (**b**) Peak 24 on chr3

containing *RUNX2*. For each region, bounded by dashed line in the plots, we calculated Z-transformed $F_{ST}$, between population nucleotide diversity ($d_{xy}$), nucleotide diversity ($\pi$), and Tajima's D. Each possible pairwise population contrast is shown and the genes that fall into the region of association are marked in the upper most plot. On the right, boxplots illustrate the difference between the statistic calculated in the main plot minus the median value across all regions of the genome that do not fall into regions of associations marked in admixture mapping. Values that fall above or below zero are more extreme values for that statistic compared to background values. In the boxplot, the centreline indicates the median, bounded by the 25$^{th}$ and 75$^{th}$ percentile, with whiskers extending to 1.5x the interquartile range.

**Fig. S5. Manhattan plots for admixture mapping of body and beak size variation in tree and ground finches.** Top, admixture mapping comparing small, medium and large tree finches (*C. parvulus*, *C. pauper* and *C. psittacula*, respectively), classified as 0,1,2, based on beak and body size. Points in red mark those that surpass the significance threshold in the *Geospiza* admixture mapping results **(Fig. 1b**). Below, the same admixture mapping result as in Fig. 1b, for all SNPs recovered in tree finches, and the points surpassing the significance threshold set by permutation (see Methods) marked in red. The position of the locus (locus #3) containing *HMGA2* is marked in both plots.

**Fig. S6**. **Neighbour-joining trees of differentiated SNPs at the 28 admixture mapping outlier loci.** NJ trees (based on species-level allele frequencies) for each of the 28 loci using the top 100 differentiated SNPs at each locus.

**Fig. S7**. **Concordance of allele frequencies at the 28 admixture mapping outlier loci.** Concordance between each locus NJ tree (Fig. S6) to the concatenated haplotype tree (Fig. 3) on the y-axis and to the species tree based on genome-wide SNPs on the x-axis; the haplotype trees in these comparisons were based on 27 loci excluding the single locus under consideration to avoid bias. Concordance is measured as the squared correlation coefficients between the distance matrices of individual locus with the distance matrices based on the concatenated haplotype tree (Y-axis) and with the distance matrix based on genome-wide SNPs (X-axis). Distance matrices at all but two loci are more concordant with the concatenated haplotype tree than with the species tree.

**Fig. S8**. **Enrichment analysis.** The software GREAT (*3*) was used to screen for enrichment of gene annotation terms associated with the 28 differentiated regions. Annotation term sharing between significantly enriched terms from Mouse Genome Informatics (MGI) gene expression databases are shown as a heat-map indicating the degree of gene sharing between each significant category. Gene sharing was defined for category pairs, as the proportion of genes from the category with the fewest genes which also occur in the category with the highest number of genes. Annotation terms along the x-axis occur in the same order as on the y-axis and the labels on the y-axis indicate term names. Also shown for each term are -log(Bonferroni *P*-value), degree of enrichment (E) and the number of genes in each category (N).

**Fig. S9**. ***In situ* hybridization (ISH). (a)** Schematic representation of the expression pattern of *ALX1* (blue) based on ISH analysis of E6 and E7 embryos of zebra finch (*T. guttata*, n=7) and Darwin's finches (9 species, n=27: 4 *G. magnirostris*, 3 *G. fortis*, 4 *G. fuliginosa*, 2 *G. difficilis*,

2 *G. propinqua*, 2 *G. scandens*, 4 *C. psittacula*, 2 *C. parvulus* and 4 *P. crassirostris*). **(b)** mRNA expression of *ALX1* and *COL9A1* – a biomarker for craniofacial cartilage (*4*) (dark blue/purple), as detected by ISH, in mid-face longitudinal sections through the heads of zebra finch embryos. The developing beak region is shown. The expression patterns of *ALX1* and *COL9A1* are mutually exclusive, indicating that *ALX1* is active in the undifferentiated beak mesenchyme and not anymore after the onset of early chondrogenic and pre-osteogenic differentiation. **(c)** mRNA expression of *ALX1* and *COL2A1* – a biomarker for craniofacial cartilage (*4*) (dark blue/purple), as detected by ISH, in mid-face longitudinal sections through the heads of Darwin's finch embryos. The expression patterns are similar to the one in zebra finch (see panel b). The images on the bottom row represent magnified regions from the images on the top row (*ALX1*), with exception of *P. crassirostris* E7: for this embryo, an image of the *COL2A1* expression is shown. Scale bar: 250 µm.

**Fig. S10. *In situ* hybridization of *ALX1* and *RUNX2*. (a)** *ALX1* is strongly expressed in the beak region only and not anywhere else in the body, as shown in zebra finch (*T. guttata*). Scale bar: 500 µm. **(b)** Expression of *RUNX2* in zebra finch (n=2). The images on the bottom represent magnified regions from the images on the top. Scale bar: 250 µm.

**Table S1: List of all tissues used for RNA-seq for genome annotation.** The type of library preparation and kit are listed, as well as the accession numbers for raw data on the European Nucleotide Archive (ENA) database. (Excel file not included in pdf)

**Table S2: Summary of the number of samples by species and island re-sequenced and used for population genetic analysis.** For a complete list by sample, see **Data S1**.

**Supplementary Data files can be found at the following link:**
https://github.com/erikenbody/Darwins_finch_comparative_genomics/tree/master/supplementary_data_for_github

**Data S1:** List of all re-sequenced samples included in this study, species, island, sequencing depth, NCBI BioProject ID, and SRA accession numbers.
**Data S2:** Divergence statistics for all 28 loci identified in association analysis. The protocols for identifying association regions and calculation of divergence statistics are presented in Methods. Median values are reported for all 20 kb windows within the association for $F_{ST}$, $d_{xy}$, $d_a$, and time since divergence. A list of genes highlighted in the GREAT analysis are followed by a column containing all genes identified within the region using the annotation pipeline reported in Methods. Ensemble gene IDs are the final column.
**Data S3:** Diversity and divergence plots as described for **Fig. S4**, for all 28 loci.
**Data S4:** Heatmaps for the most highly differentiated SNPs at all 28 loci. Per-species average allele frequencies for the SNPs reaching statistical significance in the ground finch admixture mapping are presented. The frequencies of the major allele defined in the small ground finch (*G. fuliginosa*) are shown for 18 species of Darwin's finches, two outgroup species and birds of the Big Bird lineage. The data are summarized in **Fig. 2**.
**Data S5:** Genome-wide visualization of the fraction of introgression statistic $d_{f.}$ $d_{xy}$ genetic divergence in each of the divergence loci between *G. magnirostris* and *G. propinqua*, as well as the $d_f$ values from **Fig. 3** and the heatmap from **Fig. 2.**

**Data S6:** Description of genes highlighted in **Fig. 4b**

#### Supplementary Methods

**Sample collection.** Blood samples were collected from various Galápagos islands as part of sampling described elsewhere (*2, 5*) and stored on EDTA-soaked filter paper in Drierite to preserve red blood cells for DNA extraction later. This included 101 individuals of 8 different species (**Data S1**). An additional 18 samples of 6 species were captured using mist-nests on San Cristóbal in 2018 for this study and re-sequenced using short-reads (**Data S1**). In total (combined with sequences from previous studies), our sampling includes all 18 species of Darwin's finches, the hybrid Big Bird lineage, and two outgroup species that sum to a combined sampling of 321 individuals (**Data S1**). Out of the individuals captured on San Cristóbal in 2018, one small tree finch was sampled for targeted long-read sequencing (see below). Sampling was conducted in accordance with protocols of Princeton University's Animal Welfare Committee. Embryos for RNA-seq for annotation ($n$ = 11, 3 species) and for differential expression ($n$ = 9, 6 species) analyses, and for *in situ*-hybridization ($n$ = 27, 9 species), were collected on Santa Cruz, Genovesa, and Pinta as described (*6*) (**Table S1**). Embryos were stored in methanol or RNA*later* (ThermoFisher, CA) until further use.

**ONT sequencing.** Pilot experiments prior to the expedition indicated that avian DNA (chicken and Darwin's finches) was not sequenced efficiently. We decided to develop a protocol to optimize yield from avian DNA sequencing with the MinION and hypothesized that the issue, was partially due to avian DNA being enriched in molecules carrying certain motifs or creating certain types of secondary structures which promoted blocking of nanopores and thus resulted in premature loss of actively sequencing nanopores and ultimately poor sequencing yields. We further hypothesized that T7 endonuclease I treatment of DNA prior to library generation would increase yields, because T7 endonuclease I has been used to increase sequencing yield in an ONT protocol for sequencing DNA amplified using Bacteriophage Φ29 polymerase, and has a broad substrate specificity, involving four-way junctions, various branched structures and single-base mismatched heteroduplexes (*7*). We first used g-TUBEs (Covaris) to achieve a controlled fragmentation of the isolated DNA to approximately 6-10 kb. Electrophoresis of the original g-TUBE fragmented DNA side-by-side with the T7 endonuclease I treated sample revealed a low molecular smear unique to the treated sample. In order to efficiently remove DNA cut by T7 endonuclease I, we used a custom SPRI-bead mixture (SeraMag SPRI beads, Thermo) capable of retaining DNA above the size of approximately 4-5 kb, thus selecting against the DNA cut by T7 endonuclease I. We employed the T7 endonuclease I cleavage protocol to prepare DNA for library generation for libraries aimed to generate reads in the 6-10 kb size range. For libraries where we aimed at sequencing longer molecules, we did not conduct any T7 cleavage. In the case of T7 cleavage of DNA prior to library generation we ran agarose gel electrophoresis before advancing to library isolation to ascertain successful removal of the low molecular weight DNA smear by the SeraMag speed bead mixture.

DNA was isolated by two main protocols, depending on whether we aimed to construct DNA libraries in the 6-10, 20-30 or 50+ kb size ranges. For all samples we started out with small amounts of blood (5-20 $\mu$l) sampled from wing veins of birds using glass capillaries. Immediately following sampling, the blood was deposited in a tube containing 1 ml cold PBS (10 mM EDTA). The sampled blood was kept cold in a Styrofoam box containing ice packs until back at the laboratory facility. Nuclei were isolated by adapting the protocol for cell culture in the Qiagen Genomic DNA Handbook, using ice cold buffer C1 (Qiagen). Following isolation of nuclei, we used either Blood and Tissue spin columns (Qiagen) or NaCl/ethanol precipitation to isolate <30kb or HMW DNA, respectively. Regardless of means of DNA isolation we included a DNA cleanup step using SPRI-beads prior to library generation. SeraMag SPRI-beads were prepared by mixing 10 ml of 5M NaCl, 500 $\mu$l 1M tris, 100 $\mu$l 0.5 M EDTA and 10 ml $H_2O$ was mixed together in a 50 ml Falcon tube. One ml SeraMag beads were washed 5x in 1 ml TE-buffer using a magnet stand and the final 1 ml volume was added to the 20.6 ml solution. A 50% weight/volume solution of PEG8000 in water was prepared and 18 ml of this mix was added to the 21.6 ml bead solution. The content was rigorously vortexed. 27.5 $\mu$l of Tween-20 was added to the mixture and the volume was adjusted to 50 ml with $H_2O$. The final mixture was vortexed and then aliquoted to 1.5 ml Eppendorf tubes. The performance of the bead mix at different bead/sample volume ratios was evaluated using a mixture of Bacteriophage lambda DNA and 1 kb DNA ladder.

From one small tree finch (*C. parvulus*) individual (STF5) we produced four LSK-108 libraries from g-TUBE fragmented DNA and five LSK-108 libraries made from High Molecular Weight DNA, using the protocol *Genome sequencing by ligation*, *selecting for long reads* (Oxford Nanopore Technologies). The libraries were sequenced on a GridION instrument (Oxford Nanopore Technologies) according to the manufacturer's instructions. Altogether, from individual STF5, we generated 32.3 Gbp of sequence data, corresponding to approximately 30x coverage.

**HiC analysis.** We were not able to perform HiC analysis at the Galápagos Science Center but were able to export fresh blood from a *G. fortis* individual and therefore the HiC analysis was based on a *G. fortis* individual although the long read data were from a *C. parvulus* individual. The blood was kept refrigerated until departure from Galápagos and was kept chilled (2-10$^{o}$C) until arrival at Uppsala University, where a nuclei isolation protocol (Buffer C from the Qiagen Genomics Buffer), was applied to isolate erythrocyte nuclei. Isolated nuclei were immediately snap frozen in liquid nitrogen and kept at -80$^{o}$C until further use. Nuclei were used for HiC library generation using the Arima Genomics v1 HiC kit and the resulting HiC library was sequenced on a NextSeq2000 instrument (Illumina).

**RNA sequencing.** We prepared RNA-sequencing libraries for genome annotation using a variety of library preparation protocols, tissues, and finch species. We used rRNA depletion kit from NEBNext for 6 samples of lower beak, jaw muscles, brain, gut+heart, left forelimb, and brain. An additional 4 samples were prepared using poly-A enrichment protocols from NEBNext for brain and jaw muscles tissue. Finally, a single *G. fortis* trunk sample was enriched using a Lexogon SENSE total poly-A enriched RNA-seq kit and sequenced at 3x higher depth than the remaining samples. The specific kits and sample accessions are listed in **Table S1**. All 11 libraries were multiplexed and sequenced on a flow cell of an Illumina (San Diego, CA) SP chip. For differential expression analysis, we extracted RNA from the upper beak primordia of 9 embryos from 6 species of Darwin's finches (**Table S1**), prepared cDNA libraries with the NEBNext Ultra RNA Library Prep Kit for Illumina (New England Biolabs, MA) with poly(A) selection, and sequenced them on HiSeq 4000 (Illumina, CA).

**Contrasting gene expression in beak and other tissues.** RNA-seq data were aligned to the genome assembly using STAR v2.7.2b (*8*), guided by the gtf-file from genome annotation. For each sample uniquely mapping reads for each annotated gene were extracted using the STAR option "--quantMode GeneCounts". Per sample gene counts were normalized to transcripts per kilobase million (TPM) values, first by dividing observed counts with the longest isoform length in kb, then by dividing those values with the total numbers (in millions) of uniquely mapping reads.

Obtained TPM values were used to contrast upper beak development samples (n=9) with samples corresponding to other tissues (n=7). *P*-values for differential expression from two-sided t-tests were calculated for each gene. M-values were calculated as -log2(average TPM beaks/ average TPM other tissues).

**Short-read sequencing.** 101 individuals were extracted using a custom salt preparation method (described in Enbody *et al.* (*9*)) and sequenced using the TruSeq kit (Illumina, CA). 16 additional whole-genome libraries were prepared using a custom Tn5 transposon based tagmentation protocol derived from Picelli *et al.* (*10*) and detailed in Enbody *et al.* (*9*). Briefly, we assembled the Tn5 transposon construct using the stock Tn5 (prepared by Karolinska Institutet Protein Science Facility) and the following primers (*10*):

Tn5MErev: 5;-[phos]CTG TCTCTTATACACATCT-3’
Tn5ME-A (Illumina FC-121-1030): 5’-TCGTCGGCAGCGTCAGATGTGTATAAGAGACAG-3’
Tn5ME- B (Illumina FC-121-1031): 5’-GTCTCGTGGGCTCGGAGATGTGTA TAAGAGACAG-3’

***In situ* hybridization.** Tissue sectioning and fixation, probe hybridization, and signal detection were performed according to previously published protocols (*11*) with probe concentration of 0.5 ng/µl. DIG-labelled antisense riboprobes were generated by PCR followed by RNA synthesis according to standard procedures, using T3/T7 promoter primers combined with the following gene-specific primers: *ALX1* F: 5’-CAGGACAGCAACGTCAACTA-3’, R: 5’-AAGCCTGTGTAGCCAGAATC-3’ (581 bp); *COL2A1* F: 5’-GCAAGGCCAAGGAGAAGAA-3’, R: 5’-TGATTCTGGTGTTGGGATGAG-3’ (683 bp); *COL9A1* F: 5’-CTGGCCCAAAGGGTAATAGAG-3’, R: 5’-ACCAAATTCTGGCCTCCTAAG-3’ (608 bp); *RUNX2* F: 5’-GAACCAGGTGGCCAGATTTA-3’, R: 5’-GACTGGCGGTGTATAGGTAAAG-3’ (619 bp).

**Genome assembly.** We basecalled the raw sequencing data using Guppy v. 1.6 (ONT) and further processed the basecalled data by adapter removal and splitting of reads with internal ONT adapters using the tool Downpore (https://github.com/jteutenberg/downpore). Adapter trimmed reads from one individual (STF5) were subjected to contig assembly using wtdbg2 version 2.5 (*12*) using the following parameters (-p 21 -AS 2 -s 0.05 -L 5000 -g1.1g). The resulting contig assembly featured 1,980 contigs and this assembly was polished first with two iterations of racon (v1.3.0), then with Medaka v.0.10.0 (ONT), and lastly by pilon v1.22 (*13*) using LongRanger (v2.2.2, 10X Genomics) alignments of reads from a Chromium Genome library (10X Genomics). In order to assure the contig assembly did not have problems associated with retention of both copies of divergent haplotypes we aligned a 90x Illumina data from the same individual against the contig assembly and searched for duplicated sequence which would be visible as a bimodal distribution of depth of coverage. As we could not see any signs that the assembly was partially diploid we concluded that the assembly was not in need of haplotig purging. Scaffolding was done using HiC data from a *G. fortis* male (Arima Genomics v1 kit). Manual curation of the 3D-DNA scaffolded genome was performed using the software Juicebox (*14*) where we used alignments of the scaffolded assembly against both the linked-read assembly generated using Supernova2 (10X Genomics) and the zebra finch reference genome to guide us in cases where it was difficult to infer the correct contig order from HiC contact intensities. In this curation process we required both the Supernova2 assembly and the zebra finch assembly to be in disagreement with the HiC scaffolding for manual curation of a contig orientation to be triggered. Chromosome naming was based on syntenic comparison with the zebra finch genome assembly. We used BUSCO v.4.06 (*15*) to evaluate the presence of conserved orthologs in the scaffolded genome assembly for using the `aves_odb10` database which reported 96.1% complete gene models and 1.2% fragmented gene models (i.e. 97.1% of all conserved avian genes are present in the reference assembly).

**Gene and repeat annotation.** We used trim-galore (v0.4.4, https://www.bioinformatics.babraham.ac.uk/projects/trim_galore/) with the settings (--length 36 -q 5 --stringency 1 -e 0.1) to trim RNAseq reads. Trimmed sequences were provided to Ensembl (U.K.) for genome annotation. The resulting gene predictions were blasted against the UniProt high quality protein database (*16*) to infer orthologous gene names. We used Repeatmasker (v4.0.8) (*17*) using the -q (`quick`) option and the custom repeat library for Aves to determine repeat content in the genome.

**Short-read variant analysis.** We mapped short-reads for 321 individuals to the Camarhynchus_parvulus_V1.1 genome assembly (GCA_902806625.1) using BWA mem v0.7.17 as included in the Sentieon tools wrapper (*18*). Next, variants were called using a modified version of Haplotype Caller (GATK 4.1) (*19*) included within Sentieon tools, and joint genotyped using the GVCFtyper module of Sentieon tools which is a wrapper around GenotypeGVCFs from GAKT 4.1. We ran GVCFtyper with the --emit_mode CONFIDENT to output high confidence invariant sites in addition to variant sites. We applied the following filters using VariantFiltration (GATK v4.1.4.1) to SNPs in the callset:
QD < 2.0, FS > 60.0, MQ < 40.0, MQRankSum < -12.5, ReadPosRankSum < -8.0, SOR > 3.0

In addition to genotype filters for:
DP < 2, DP > 100, GQ < 20

We set filtered genotype calls to no call using Select Variants (GATK, v4.1.4.1) then removed sites not passing filters using bcftools (http://www.htslib.org/, v1.10). We created a separate file of high confidence invariant sites requiring <50% missing data using vcftools v0.1.16 (*20*). The resulting invariant file was concatenated with the high-quality SNP set using bcftools concat.

**Genotype phasing.** We inferred haplotypes using a combination of WhatsHap (*21*) (v0.18) and SHAPEIT4 (*22*) (v4.1.3) to generate a phased variant callset of biallelic SNPs. We used WhatsHap under default settings to identify phase informative reads (i.e., a read spanning at least two heterozygous sites) and the resulting VCF file was used as input for SHAPEIT4. WhatsHap was run separately for each individual, after which all individual VCFs were merged using the bcftools merge command. We ran SHAPEIT4 with a constant recombination rate of 1 cM per Mb and an expected error rate in the phase informative read sets of 0.0001 (the default setting).

**Construction of linkage map.** We computed recombination rate (rho) for 25 samples of *C. parvulus* from five islands using LDhelmet v1.10 (*23*). We additionally used three *T. bicolor* and five *L. noctis* samples from Barbados as outgroups. The phased haplotypes generated using SHAPEIT4 were converted to the required genotype format of LDhelmet using the vcftools –ldhelmet flag. In order to identify the ancestral state, we selected at each site the allele present in >60% of outgroup samples and following Singhal *et al*. (*24*) we assigned prior probabilities of 0.91 for the ancestral base and 0.03 of the remaining bases in order to account for the possibility of incorrect ancestral inference. At unresolved sites (e.g., missing data in outgroup samples), we used a prior of the stationary distribution of allele frequencies from the mutation rate matrix following the LDhelmet (*23*) method (also see Cam_parv_mutation_matrix.xlsx). Using genome-wide sites, we used the LDhelmet *find_confs* module with a window size of 50 SNPs to generate a haplotype configuration file. We next generated a likelihood lookup table using the *table_gen* command using the recommended rho grid values of (-r 0.0 0.1 10.0 1.0 100.0) and a previously published Watterson's theta estimate of 0.00126(*2*). Last, we used the *pade* command to generate Padé coefficient tables using the same estimate of Watterson's theta and the suggested 11 coefficients. We ran LDhelmet in 50 bp windows for 10,000,000 iterations (discarding the first 200,000 as burn-in), and the following additional flags:

--max_lk_end 100 --prior_rate 0.05 -b 5.0

A previous study identified a block penalty of 5, as used here, as the ideal value for demonstrating fine scale recombination patterns (*24*). We extracted ρ/bp, representing the population scaled recombination rate, using *post_to_text* within LDhelmet. We summarized ρ by 20 kb sliding, non-overlapping windows (using the R package windowscanr, https://github.com/tavareshugo/WindowScanR), and report ρ/kb in comparison with diversity statistics. We additionally converted ρ to recombination fraction following r = ρ/2cNe (*1*) and Ne = 33,861 (*2*). Recombination fractions were converted to cM/Mb using the Morgan function r * 100 (for all plots in **Fig. S2**).

**Admixture mapping.** We used GEMMA (*25*) (genome-wide efficient mixed-model association 0.98.1, 2018-12-10) to search for loci associated with the scaling up in beak and body size from *G. fuliginosa* (*n* = 26), *G. fortis* (*n* = 34) and *G. magnirostris* (*n* = 15) by recoding each species as 0,1,2, respectively, and estimating the strength of association for each SNP under an additive model. In order to account for relatedness among samples, we first calculated a relatedness matrix. As input for the relatedness matrix, we pruned autosomal SNPs by retaining only 1 randomly selected SNP per 20 kb in bcftools +prune (v1.11-54-gaf54707) to reduce the effect of linkage disequilibrium on the relatedness calculation. We converted the resulting pruned VCF file to bimbam_dosage format using qctool (https://www.well.ox.ac.uk/~gav/qctool_v2/) and calculated the relatedness matrix using the GEMMA flag -gk 1. A separate relatedness matrix was calculated for SNPs located on chromosome Z using identical parameters. We ran linear mixed-effects models (LMM, -lmm 2 option in GEMMA) with a maximum likelihood estimation (MLE) using the relatedness matrix calculated previously as a covariate and separately for all autosomes and chromosome Z. The same analysis was run for small, medium and large tree finches (*C. parvulus n* = 25, *C. pauper n* = 10, *C. psittacula n* = 11). For *Geospiza*, we used the code implemented in gemma-wrapper to run permutations on 100 independent runs of shuffled phenotype values to establish a baseline significance threshold for SNPs exceeding the $67^{th}$ percentile of the distribution of *P*-values. This resulted in thresholds of $-\log_{10}$(*P*-value) = 7.3 (*P*-value < $5 \times 10^{-8}$). Using custom R scripts (v4.0.3) (*26*), we identified genomic regions that contain SNPs exceeding this threshold by extending peaks from the top SNP of association down to an arbitrary $\log_{10}$(*P*-value) threshold of 3.6. Within these regions, we merged all SNPs within 75 kb into a single genomic interval using the GenomicRanges package and requiring each interval to have more than 1 SNP. This resulted in 28 genomic regions of association on autosomes that exceeded the threshold set by permutation. In order to extract SNPs tagging the haplotypes of association, we selected the top SNPs of each region use the `slice` function in the R package dplyr to choose the top 100 SNPs based on their strength of association. In some genomic regions, many SNPs are linked and those with identical *P*-values were retained for analyses including the top SNPs, meaning that in 15 genomic regions shown in **Data S4** include > 100 SNPs. In four regions, <100 SNPs exceeded the thresholds defined here.

**Genetic diversity and divergence analyses.** We calculated unbiased estimates of $F_{ST}$, $d_{xy}$ and $\pi$ using the software pixy (v1.0.0.beta1) (*27*). For this analysis, we used the VCF that included both high confidence invariant sites and biallelic SNPs (see methods under Short-read variant analysis). We calculated $F_{ST}$ and $d_{xy}$ for all three possible combinations of *G. fuliginosa* ($n$ = 26), *G. fortis* ($n$ = 34) and *G. magnirostris* ($n$ = 15) and nucleotide diversity (π) for each species in non-overlapping 20 kb windows and by applying a minor-allele frequency filter of 0.05 within the pixy command module. We additionally removed windows with < 100 SNPs. Time to divergence was estimated by the conversion of genetic divergence ($d_a$) to time using $T = d_a/(2\mu)$ and a mutation rate of 2.04 X $10^{-9}$ (*2*). Corrected estimate of sequence divergence ($d_a$) was calculated as $d_{xy}$ – average nucleotide diversity in both populations of the comparison (i.e., mean genome-wide π in *G. fuliginosa* and *G. magnirostris*).

Tajima's D was calculated using the same input for 20 kb genomic windows using vcftools (v0.1.16) (*20*). In order to overlap divergence statistics, which were calculated in windows, with per-SNP admixture mapping results, we used the "windowscanr" R package (Hugo Tavares (2021). windowscanr: Apply functions using sliding windows. R package version 0.1.) to calculate mean $-\log_{10}$(*P*-value) in 20 kb windows and overlap these values with the divergence statistics using the "fuzzyjoin" R package (David Robinson (2020). fuzzyjoin: Join Tables Together on Inexact Matching. R package version 0.1.6. https://CRAN.R-project.org/package=fuzzyjoin) with a max distance of 10 bp. We calculated correlation coefficients between $F_{ST}$ and $-\log_{10}$(*P*-value) using the base R command "cor" and report $R^2$ as the square of this output. We calculated $R^2$ between recombination rate ($r$) and nucleotide diversity using the same method.

**Analysis of introgression.** We first used the software Dsuite v0.4 (*28*) to calculate ABBA and BABA sites for all species and islands of *G. magnirostris*, *G. propinqua*, and *G. scandens* using *T. bicolor* and *L. noctis* as outgroups to determine the derived state for each allele. We selected these three ingroup taxa, because *G. scandens* and *G. propinqua* share a most recent common ancestor more recently than either does to *G. magnirostris(2).* We first established that *G. propinqua,* which is only found on Genovesa and sister to the allopatric common cactus finch *G. scandens*, shares the largest number of derived alleles with the population of *G. scandens* on Pinta. We subsequently used *G. scandens* samples from Pinta as P1 in a phylogenetic arrangement as shown below:

(P1, P2)(P3) = (*G. scandens* - Pinta, *G. propinqua* - Genovesa)(*G. magnirostris* - Genovesa)

The ABBA and BABBA counts and associated D statistics are as follows:

BBAA = 451573
ABBA = 412104

BABA = 348466
D = 0.084
Z = 8.23
$P = 9.0 \times 10^{-17}$

We next used Dsuite to calculate the fraction of introgression in sliding windows of 50 SNPs (by 25 SNPs) across the genome as measured by the $d_f$ which is distinct from other measures of introgression in that it accounts for divergence between each comparison in the trio, to better assess the difference between ILS and introgression (*29*).

We additionally compared $d_{xy}$ values in the 28 regions between *G. propinqua* and *G. magnirostris* using the same methodology in Pixy as described above.

**Gene annotation enrichment.** The software GREAT v3.0 (*3*) was used for annotation term enrichment analysis. Significant loci from admixture mapping were converted to chicken genome assembly (Galgal6, NCBI accession GCA_000002315.5) coordinates using Progressive Cactus v0.1-c4bed56(*30*) to first perform whole genome alignment between the *C. parvulus* genome assembly and Galgal6. Next, halLiftover from the halTools package(*31*) was used to obtain the corresponding chicken coordinates. We then used the UCSC chicken to human chain file to extract syntenic human (hg19) coordinates. Extracted human coordinates were merged and then submitted to the online GREAT tool v3.0 (http://great.stanford.edu/public/html/) using standard settings. Enriched terms (Bonferroni *P*-values < 0.05) from hypergeometric tests were retained and, as a conservative measure, only terms showing enrichment values ≥2 were retained for further analyses and plots.

**Phylogenetic reconstruction.** Neighbor-joining (NJ) trees were generated by first calculating per-species average allele frequencies from genotypes recorded in VCF files, then reformatting allele frequencies to serve as input for PopTree (*32*), where NJ trees were generated using the "poptree" command with $F_{ST}$ as the distance option and 100 bootstrap iterations. For the genome-wide species tree the input VCF was filtered to include all SNPs with a minor allele frequency ≥0.15. The topology of this tree matches previous studies (*2*). For the tree based on the top 100 SNPs from each of the 28 loci under selection, the top SNPs from each locus were combined in a single PopTree input file. In order to not bias the tree inference from the regions with >100 top SNPs (see Admixture mapping methods), we reduced the number of SNPs to exactly 100 SNPs for all regions (except those where < 100 surpassed the threshold) by sorting SNPs first by $-\log_{10}$(*P*-value) as before, then choosing randomly among positions that include ties and overlap rank 100 and above. NJ trees were exported to Newick format using the "cnvtre" command within PopTree. To analyse how concordant trees based on the 100 SNPs from each of the 28 loci under selection were compared to (1) the combined tree based on Top SNP for all 28 loci and (2) to the tree based on all genome-wide SNPs with a minor allele frequency ≥0.15 we exported genetic distance matrices for each tree from PopTree (*32*). Pearson Product-Moment Correlation Coefficients were then determined for each genetic distance matrix pair.

We generated a "cophylo" plot to compare topologies between the species tree and the haplotype tree generated for the concatenated SNP matrix of all 28 loci using the R package phytools v0.7 (*33*). Newick trees were loaded into R with the R package ape v5.5 (*34*) and a chronogram generated for the species tree using the phytools commands makeChronosCalib and chronos, setting a maximum age at 1 million years following Lamichhaney *et al.* (*2*). Next, we ran the cophylo command with default settings (include rotate = T) which attempts to optimize the orientation of tips in both trees.

Separately, we created a maximum likelihood phylogenetic tree for all *G. fuliginosa*, *G. fortis* and *G. magnirostris* samples included in the admixture mapping analysis (**Fig. S3)**. For this analysis, we used the pruned genotype data that was used for generating the relatedness matrix in GEMMA (see Methods above). We used the BEDtools (v2.29.2) (*35*) intersect command with the `-v` flag (while supplying a bed file of genomic regions) to retain only genomic regions laying outside the 28 loci identified in admixture mapping and converted the VCF file to fasta format using vcf2phylip v2.4 (*36*). We included a randomly selected allele at each heterozygous position by supplying the –resolve-IUPAC flag in vcf2phylip. We ran FastTree (*37*) to generate a maximum likelihood phylogenetic reconstruction with the following settings:

```
FastTree -nt -log finch_fasttree_log_file -gtr -fastest
finches_geospiza.fasta
```

We used plink2 (*38*) (v2.00a2.3LM) to generate a covariance matrix of all genotypes either within or outside of the 28 genomic regions identified in the admixture mapping analysis. As input we used the pruned dataset described above that was used to generate the maximum likelihood phylogeny of the three *Geospiza* species. We also created a genotype matrix (VCF file) of only the 28 regions using a bed file of these regions supplied to BEDTools intersect. We ran the plink –pca command under default settings for both the genotype data within and outside the 28 regions of interest, and plotted the resulting covariance matrix in R (**Fig. S3b).**

**Analyses of allele frequencies.** Per-species average allele frequencies for 18 species of Darwin's finches, the Big Bird lineage and two outgroup species from genotypes recorded in VCF files were calculated using custom python scripts.

a

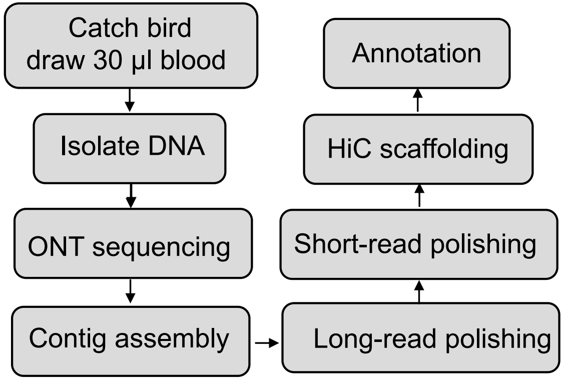


b

| | n | Length (bp) | N50 (Mb) | Gaps |
|---|---|---|---|---|
| Contig | 1,980 | 1,063,904,364 | 15.1 | 0 |
| Scaffold | 1,756 | 1,064,016,364 | 71.1 | 224 |

c

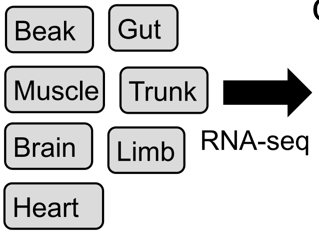


d

| Category | n |
|---|---|
| Coding genes | 16,191 |
| Non-coding genes | 883 |
| Pseudogenes | 93 |
| Gene transcripts | 25,899 |

e

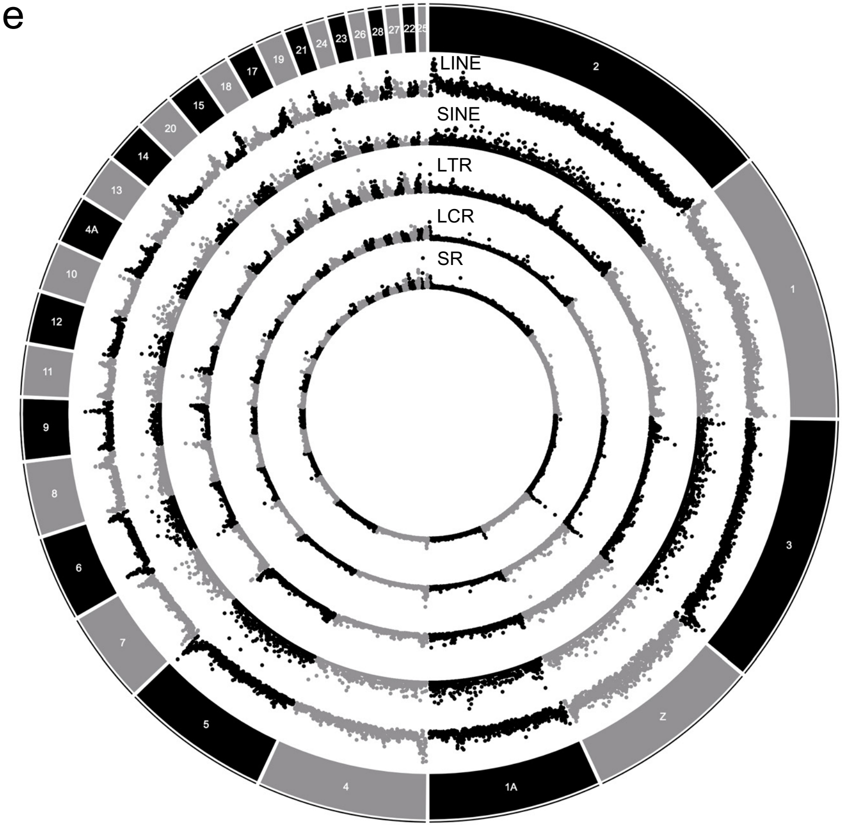


### Supplemental Fig. 1

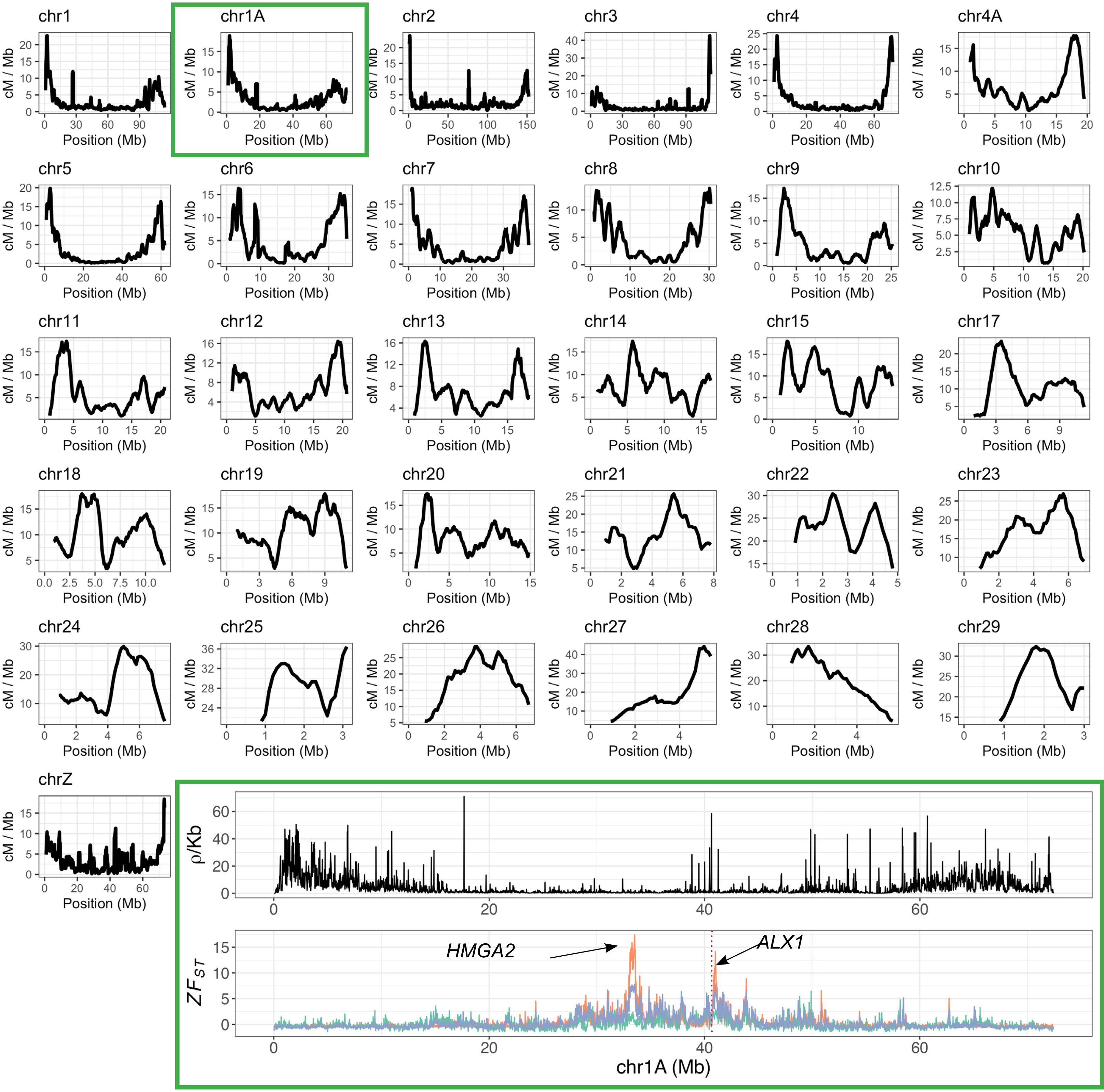


**Supplemental Fig. 2**

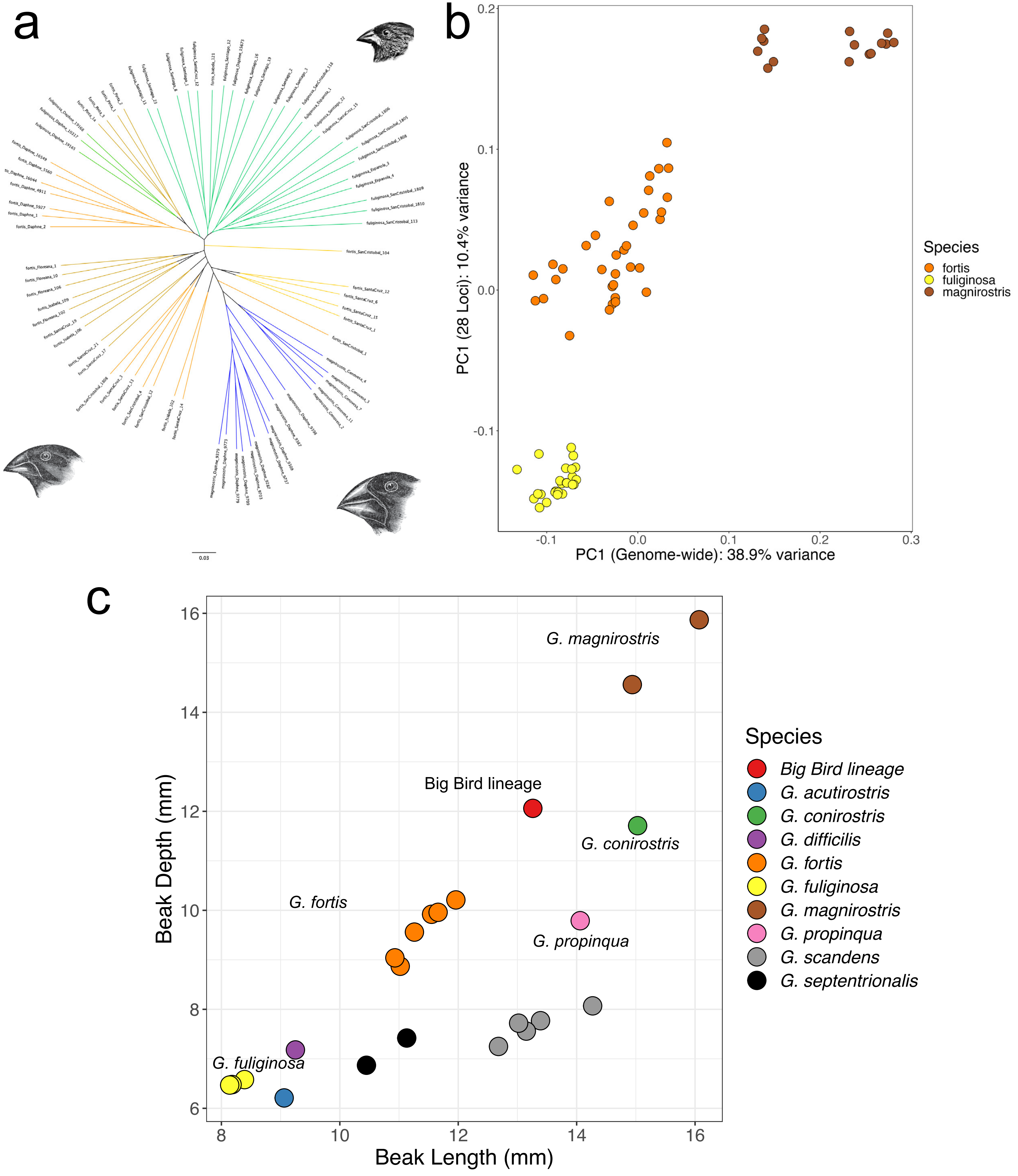


**Supplemental Fig. 3**

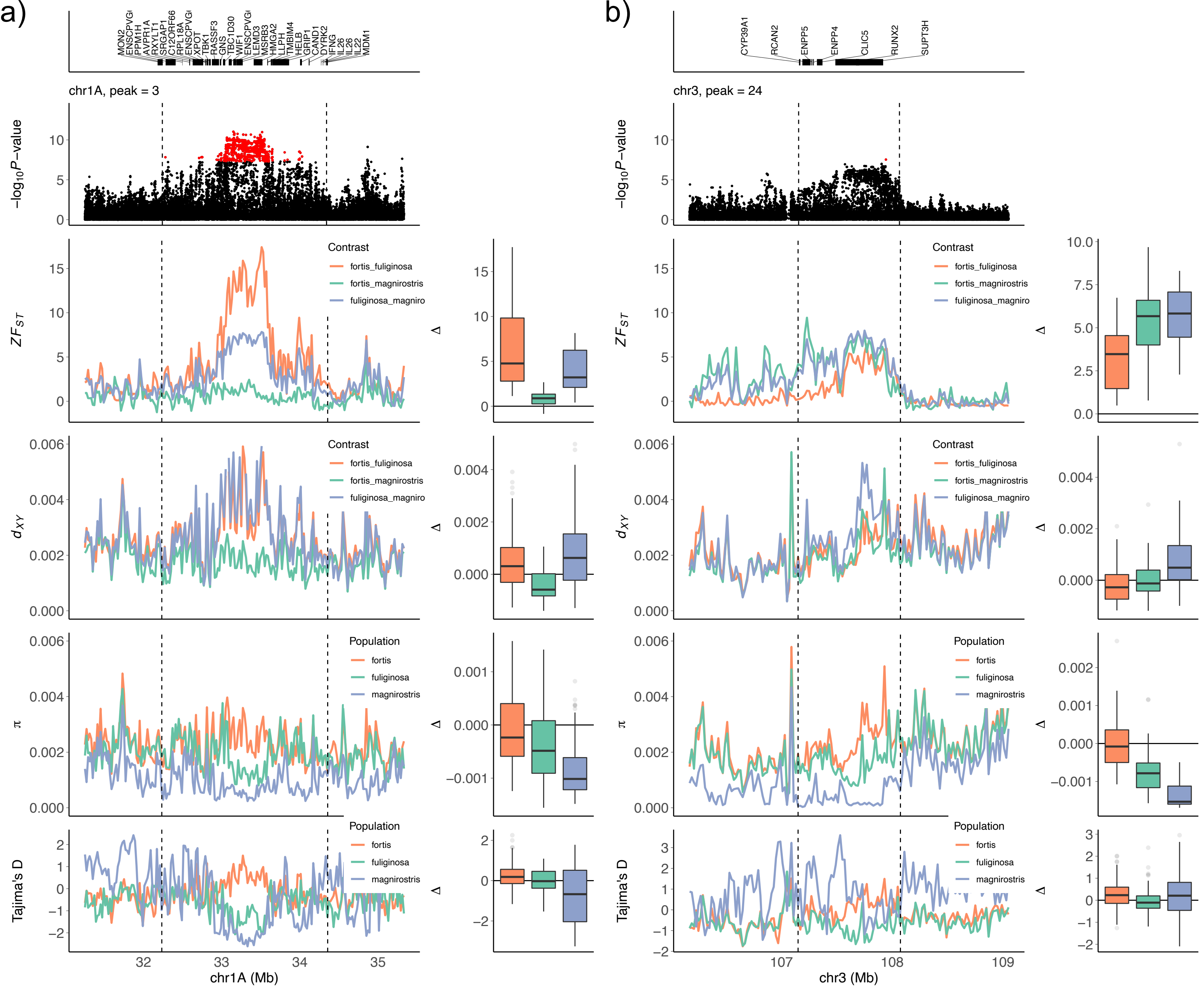


### Supplemental Fig. 4

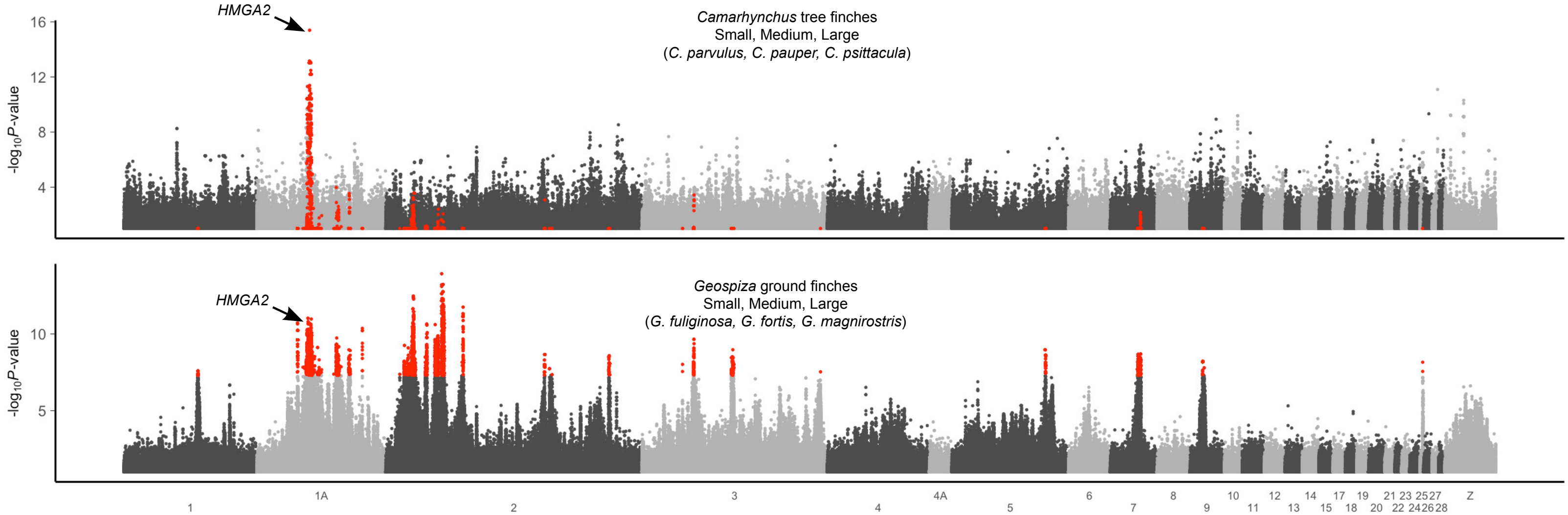


### Supplemental Fig. 5

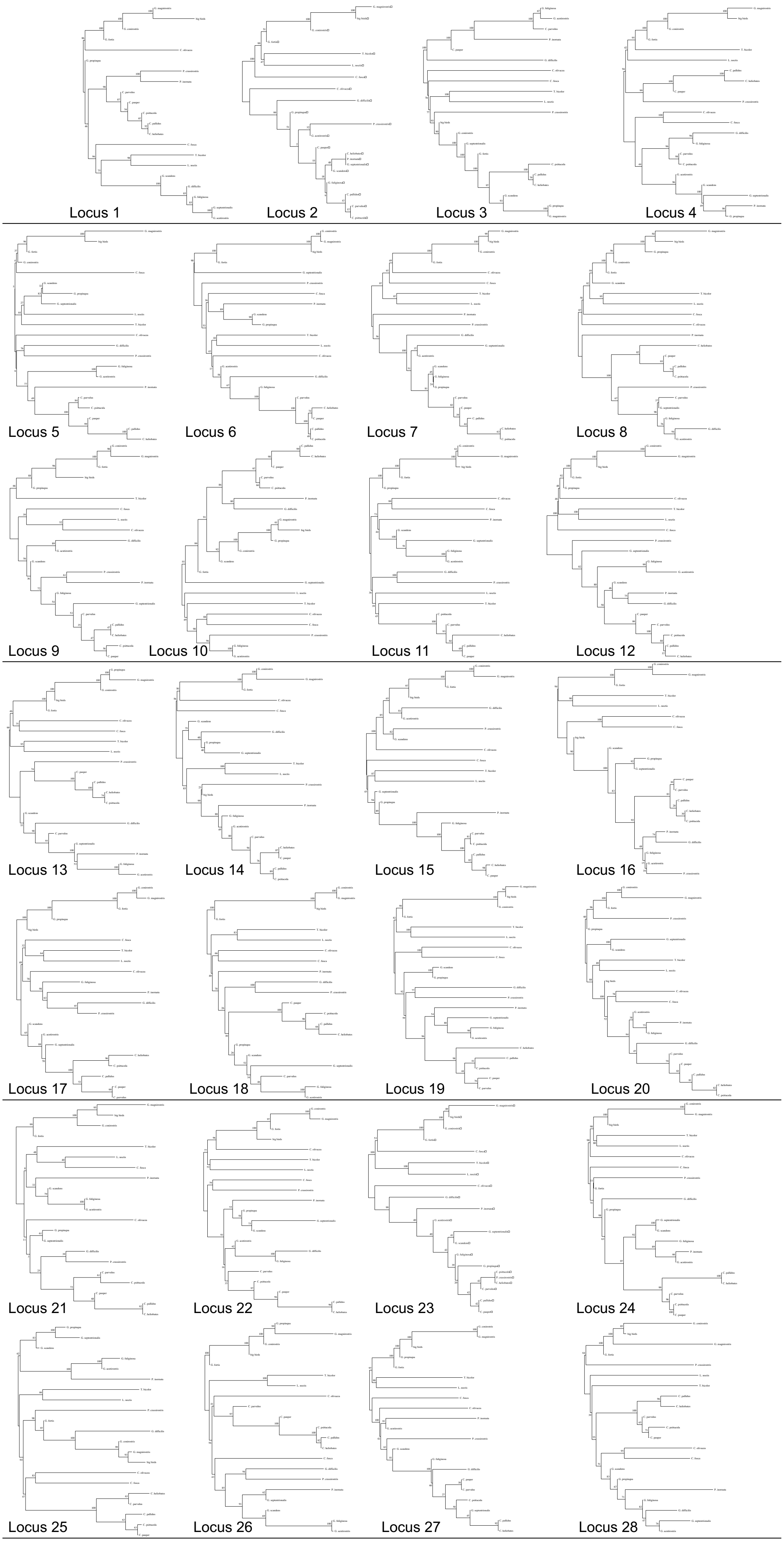


**Supplemental Fig. 6**

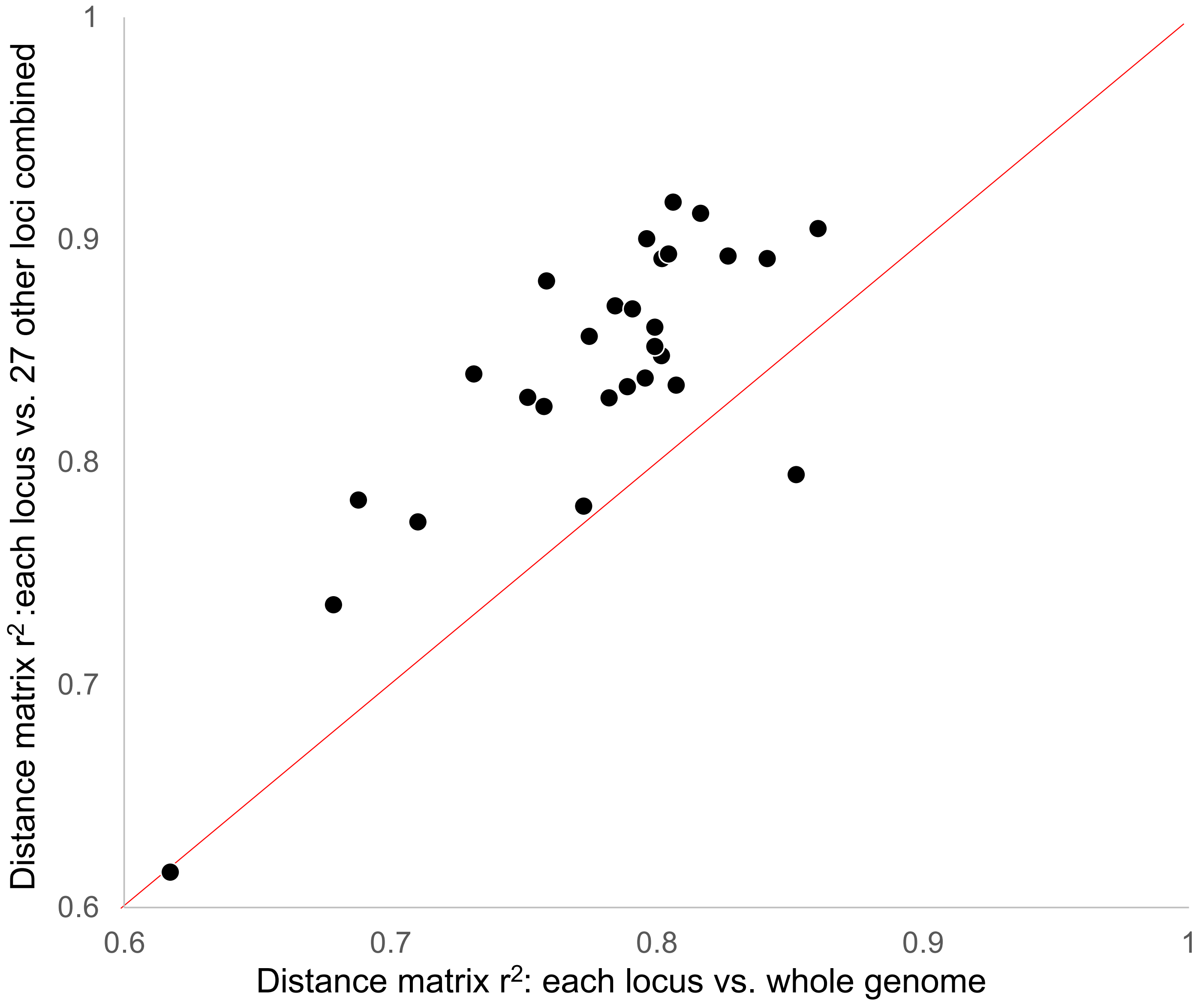


**Supplemental Fig. 7**

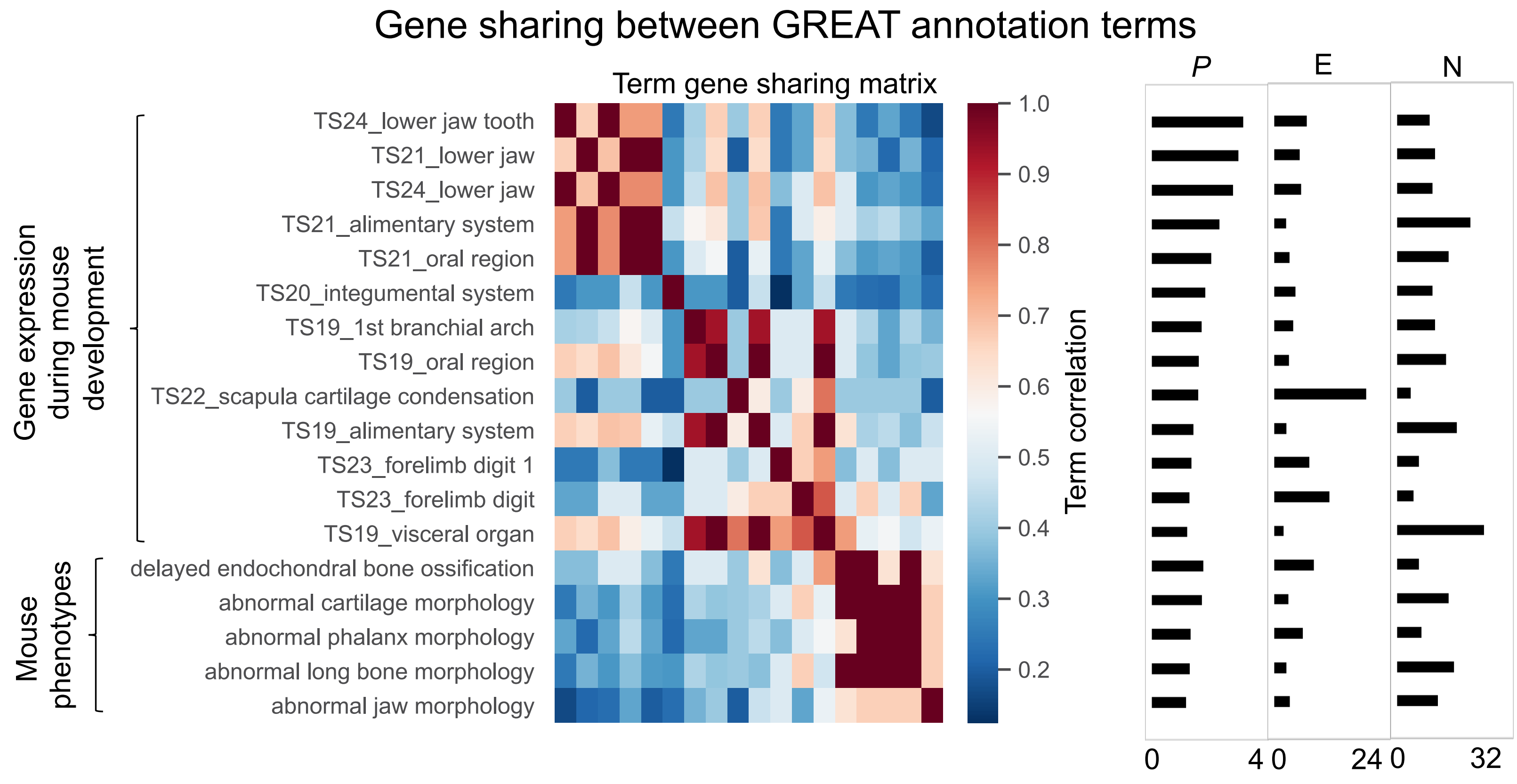


*P*= -log10(Bonferroni *P*) E= Enrichment over background N= Number of genes in category

**Supplemental Fig. 8**

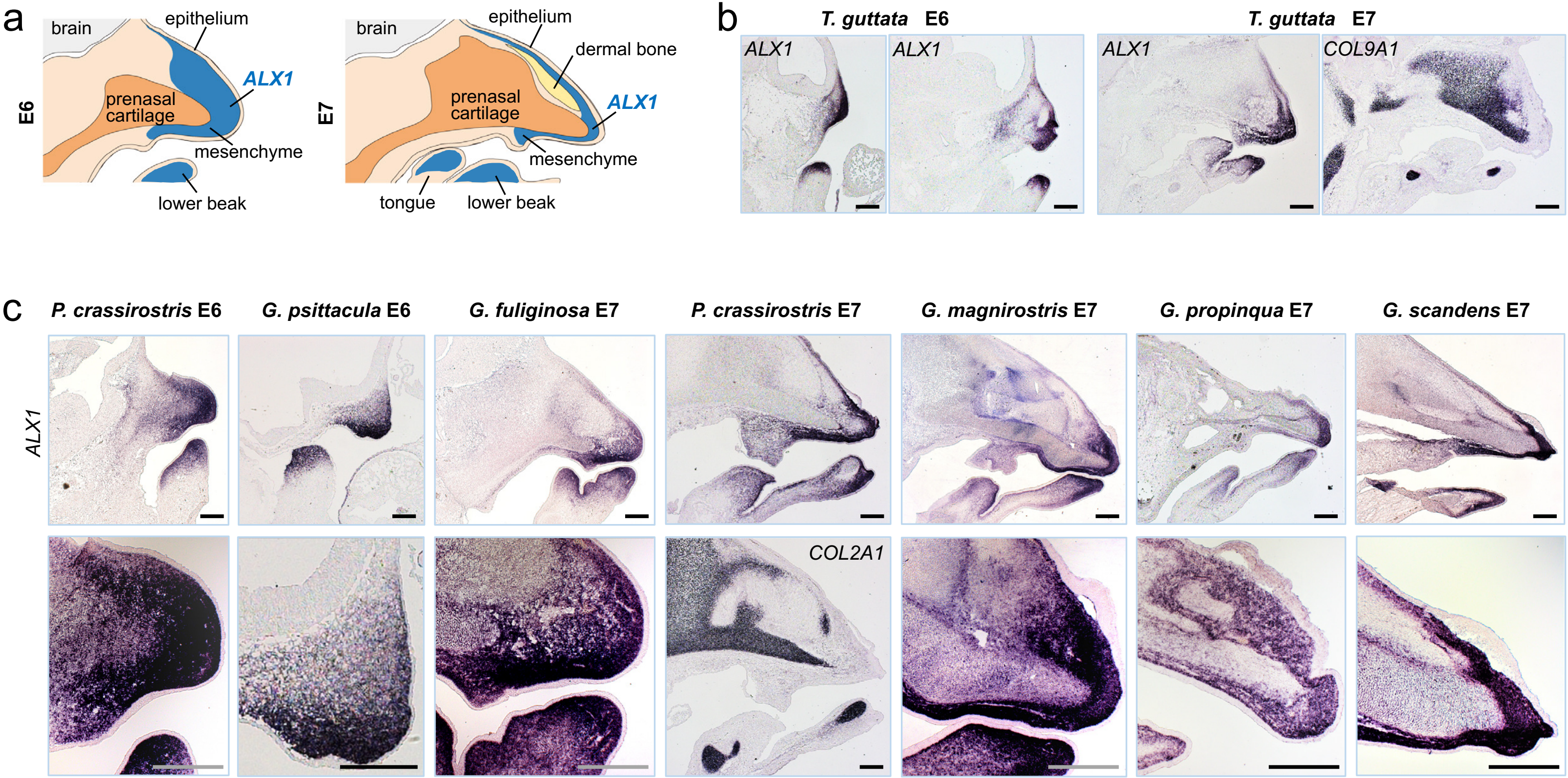


### Supplemental Fig. 9

a

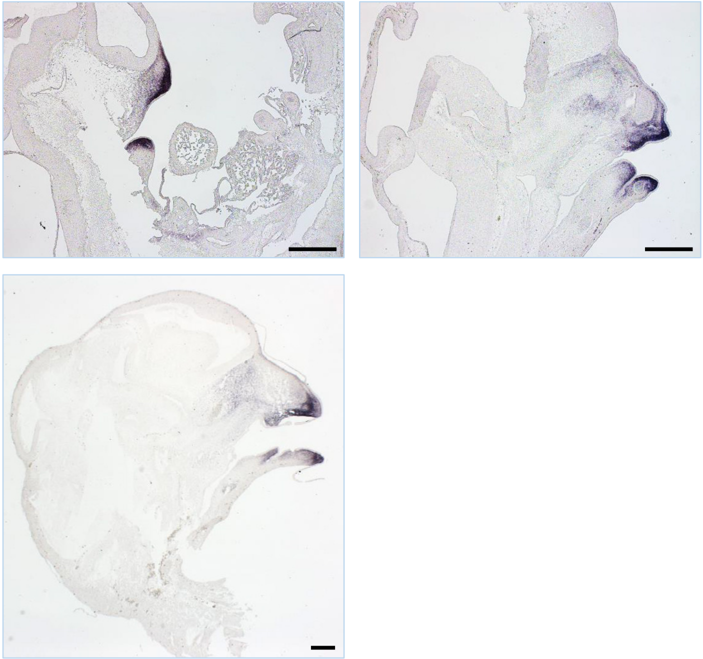


b

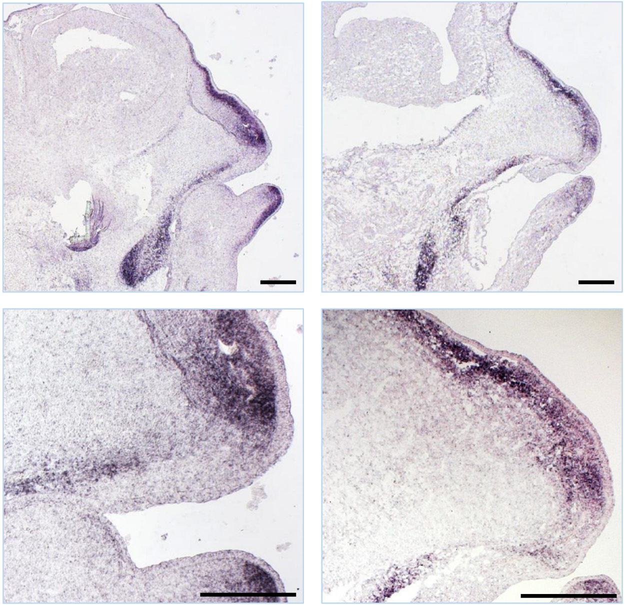


### Supplemental Fig. 10
